# Excess p50 induces Arp1-dependent dynactin clusters containing the assembly factor VezA

**DOI:** 10.64898/2026.05.29.728734

**Authors:** Jun Zhang, Rongde Qiu, Xin Xiang

## Abstract

Dynactin, a complex essential for cytoplasmic dynein function, contains an Arp1 mini-filament with a pointed-end sub-complex including Arp11 and a shoulder sub-complex including p50 and p150. We recently identified in *Aspergillus nidulans* a vezatin homolog VezA that enhances dynactin assembly, but VezA does not co-localize with dynactin, suggesting a transient interaction. It was found that overexpression of p50 separates the shoulder from the mini-filament and causes late-onset neurodegeneration in mice, but its cellular effects in the context of VezA need to be studied. Here we found astonishingly that p50 overexpression in *A. nidulans* causes Arp11, p50 (but not p150), and VezA to form co-localized clusters whose presence depends on Arp1. Formation of the VezA cluster also depends on Arp11, and VezA significantly enhances the intensity of Arp11-GFP clusters but not that of p50-GFP clusters. Moreover, an evolutionarily conserved region of p50 (aa17-25) forming a beta-sheet structure with Arp1 as revealed in previous cryo-EM studies is critical for the formation of Arp11, p50 and VezA clusters. These results suggest that excess p50 induces cluster formation driven by p50-Arp1 interactions, and that this assembly intermediate without the full shoulder traps VezA, which is involved in the Arp11-Arp1-p50 interactions.

## Introduction

The minus-end-directed microtubule motor cytoplasmic dynein-1 (called “dynein” hereafter) transports a variety of cargoes. A key dynein regulator, dynactin, is needed for the dynein-cargo interaction and cargo adapter-mediated dynein activation (McKenney et al., 2014; Olenick and Holzbaur, 2019; Rao et al., 2026; Reck-Peterson et al., 2018; Schlager et al., 2014; Yeh et al., 2012; Zhang et al., 2011; Zhang et al., 2017). Dynactin is a multi-protein complex (Schroer, 2004). Within it, the actin-related protein Arp1 forms a mini-filament of ∼36-nm (containing eight Arp1 subunits and one conventional actin subunit) that bind dynein and cargo adapters, and its two ends are capped by capping protein and a pointed-end sub-complex containing Arp11, p62, p25 and p27 (Chaaban and Carter, 2022; Chowdhury et al., 2015; d’Amico et al., 2022; Eckley et al., 1999; Gama et al., 2017; Garces et al., 1999; Grotjahn et al., 2018; Karki et al., 2000; Lau et al., 2021; Qiu et al., 2018; Schafer et al., 1994; Urnavicius et al., 2018; Urnavicius et al., 2015; Yeh et al., 2013) (Figure 1A). The rest of dynactin contains a dynein-interacting p150^Glued^ (called p150 for simplicity) that has an N-terminal microtubule-binding domain, p50 (also called dynamitin), and p24 (Echeverri et al., 1996; Gill et al., 1991; Holzbaur et al., 1991; Karki and Holzbaur, 1995; Karki et al., 1998; Okada et al., 2023; Singh et al., 2024; Vaughan and Vallee, 1995; Waterman-Storer et al., 1995). Specifically, four copies of p50, two copies of p24 and two copies of p150 C-termini form the “shoulder” of dynactin (Cheong et al., 2014; Lau et al., 2021; Maier et al., 2008; Urnavicius et al., 2015). The p50 multimer extends “tentacle”-like extended structures that bind multiple sites of the Arp1 mini-filament, which has been proposed to ensure that the Arp1 mini-filament to be assembled with a correct length of 36-nm (containing eight Arp1 subunits and one conventional actin subunit) (Urnavicius et al., 2015). A large excess of p50 has been known to separate the shoulder from the mini-filament and affect dynein function (Burkhardt et al., 1997; Echeverri et al., 1996; Eckley et al., 1999; Karki et al., 1998; LaMonte et al., 2002; Melkonian et al., 2007). Moreover, p50 overexpression in mouse neurons causes late-onset neurodegeneration similar to what happens in human amyotrophic lateral sclerosis (ALS) (LaMonte et al., 2002). Despite the clinical relevance and its significance in dynein-mediated transport of many different types of cargoes, how dynactin is assembled remains unclear.

**Figure 1.**
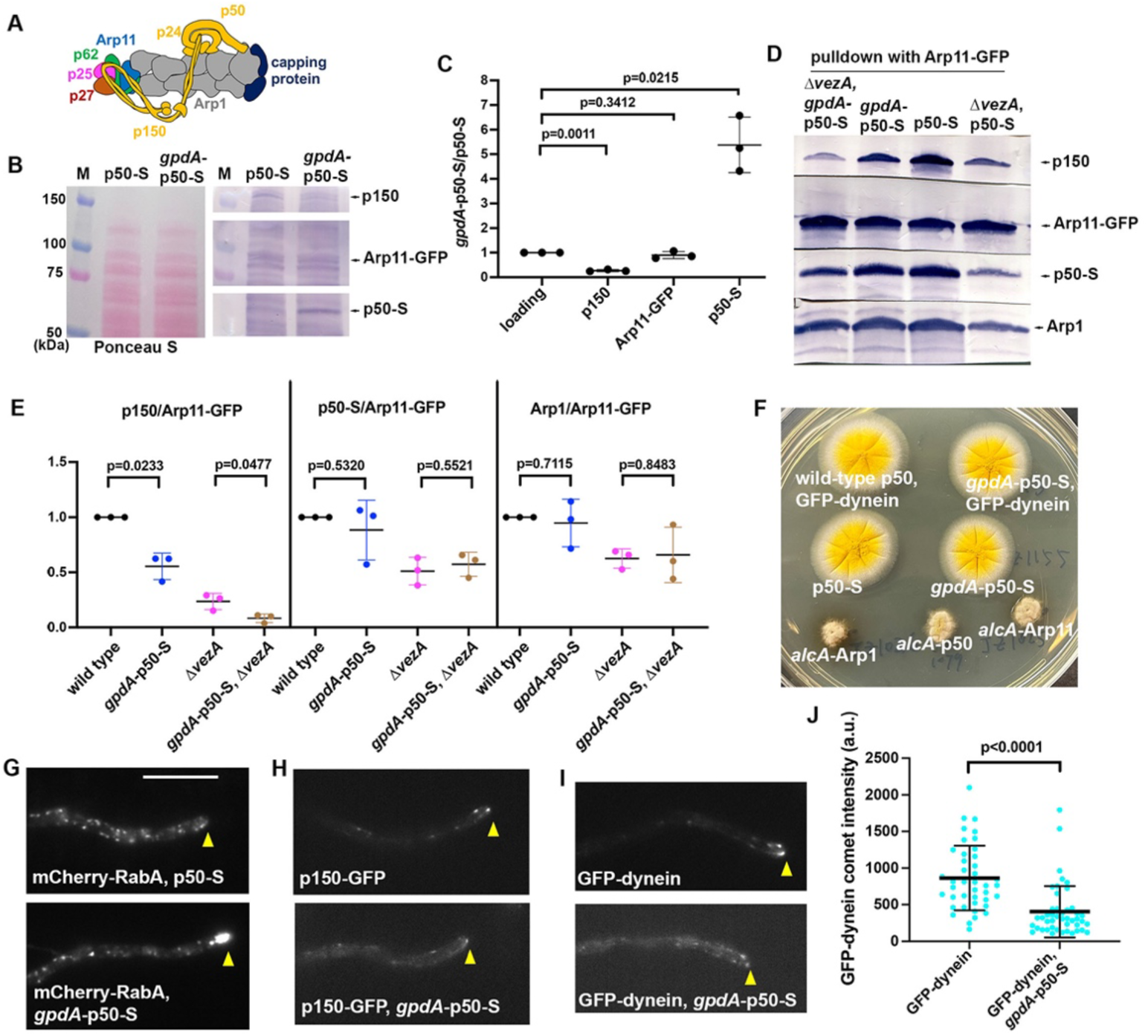
Phenotypic analysis of strains with p50 overexpression (*gpdA*-p50-S). (A) A diagram of the dynactin complex. (B) Western blots showing p150, Arp11-GFP and p50-S signals (probed with the p150, the anti-GFP and the anti-S antibodies) from the strains containing either p50-S or *gpdA*-p50-S. Ponceau S staining was also shown. M: molecular marker. (C) A quantitative analysis on the effect of p50 overexpression (*gpdA*-p50-S) on p150, Arp11-GFP and p50-S levels. The ratios of the intensity values of *gpdA*-p50-S to those of p50-S are shown for each component, and these ratios are relative to the ratio for the loading control that was set as 1. Scatter plots with mean and SD values as well as p values were generated by Prism 10 (Unpaired t tests with Welch’s correction). (D) Western blots showing p150, p50-S and Arp1 pulled down with Arp11-GFP in different strains. (E) A quantitative analysis of the effect of *gpdA*-p50-S on the amounts of p150, p50-S and Arp1 pulled down with Arp11-GFP in wild-type or Δ*vezA* background. The ratios of the intensity values of p150, p50-S or Arp1 to those of Arp11-GFP are shown. The control strain containing p50-S and Arp11-GFP is called “wild type”. Scatter plots with mean and SD values as well as p values were generated by Prism 10 (Unpaired t tests with Welch’s correction). (F) The strains with *gpdA*-p50-S form normal colony, in contrast to the conditional null mutants of dynactin that form a typical *nud*-mutant-like compact colony. (G) The *gpdA*-p50-S strain shows a defect in dynein-mediated early endosome transport as evidenced by an abnormal accumulation of mCherry-RabA near the hyphal tip. Note that >50% of hyphal-tip cells (n>100) showed this abnormal accumulation. Hyphal tip is indicated by a yellow arrowhead (same for H and I). Bar, 10 μm. (H) p150-GFP in the *gpdA*-p50-S background. (I) GFP-dynein in the *gpdA*-p50-S background. (J) A quantitative analysis of plus-end dynein accumulation. Scatter plots with mean and S.D. values were generated by Prism 10, and the p values were generated by Mann-Whitney test (unpaired).

Filamentous fungi are excellent model organisms for discovery of new proteins and mechanisms involved in long-distance intracellular transport and cellular organization (Christensen and Reck-Peterson, 2022; Driscoll et al., 2025; Geisterfer et al., 2026; Oakley and Peñalva, 2025; Riquelme et al., 2018; Rogers et al., 2026; Salogiannis et al., 2021; Steinberg et al., 2017; Vázquez-Carrada et al., 2026; Xiang and Qiu, 2020). Recently, we discovered a dynactin assembly factor VezA (vezatin homolog) in the filamentous fungus *Aspergillus nidulans* (Zhang et al., 2024). VezA was initially identified from a genetic screen as a protein important for dynein-mediated transport of early endosomes (Yao et al., 2015). Vezatin homologs in Drosophila and zebrafish are also important for dynein-mediated axonal transport of cargoes (Spinner et al., 2020). Loss of VezA in *A. nidulans* (Δ*vezA*) significantly decreases the amounts of Arp1, p50 and p150 pulled down with pointed-end proteins such as Arp11 (Zhang et al., 2024). Thus, one possibility is that VezA enhances the Arp1 mini-filament integrity. However, since the mini-filament assembly and shoulder assembly depend on each other (Zhang et al., 2024), it is not easy to pinpoint the site of VezA action. In the Δ*vezA* mutant, p150 and p50 protein levels are reduced (Zhang et al., 2024). This is most likely due to instability of proteins not bound to the complex, as p150 and p50 protein levels are dramatically reduced upon loss of Arp1, and p150 level is dramatically reduced upon loss of p50 (Haghnia et al., 2007; Minke et al., 1999; Zhang et al., 2024; Zhang et al., 2008). Based on our pulldown data, integrity of the shoulder affects that of the mini-filament (Zhang et al., 2024). This raises another possibility that VezA affects the attachment of the shoulder, which in turn affects p50-p150 stability and the integrity of the Arp1 mini-filament. Thus, although we know that VezA promotes dynactin assembly, we do not know where in the assembly pathway it acts. Moreover, although overexpressed VezA missing the transmembrane domains can pulldown dynactin components in an Arp1- and Arp11-dependent fashion (Zhang et al., 2024), the VezA-dynactin association has never been observed in live cells, possibly due to its transient nature.

In this current study, we initially sought to test if increasing the p50 level in the Δ*vezA* mutant could enhance the amount of Arp1 pulled down by Arp11-GFP, but such an effect was not observed. However, we were surprised to find that excess p50 in wild-type background induces formation of novel clusters of Arp11, p50 and VezA. We further found that these proteins co-localize in the same clusters, suggesting that p50 overexpression induces an assembly state that traps the assembly factor VezA.

## Results

### Overexpression of p50 in *A. nidulans* causes a moderate defect in dynein function and a reduction in the p150 level

To overexpress p50, we constructed a *gpdA*-p50-S allele in which S-tagged p50 is expressed from a strong promoter of *gpdA* (Pantazopoulou and Penalva, 2009; Zhang et al., 2011). To judge the extent of p50 overexpression, we used western analysis for proteins from the *gpdA*-p50-S-containing cells and a control strain with 50-S expressed from the endogenous promoter (Zhang et al., 2024). Both strains contain Arp11-GFP (Qiu et al., 2020). We found that the *gpdA*-promoter enhances the level of p50-S about five-fold (Figure 1B, 1C) and decreases the p150 protein level significantly (Figure 1B, 1C).

We then examined if *gpdA*-p50-S changes the amounts of Arp1, p50 or p150 pulled down by Arp11-GFP in wild-type or the Δ*vezA* mutant. The strains containing p50-S expressed under the endogenous p50 gene promoter were used as controls. In the wild-type background, *gpdA*-p50-S reduces the amount of p150 pulled down by Arp11 as expected since the p150 level in total extract is lower in the *gpdA*-p50-S strain (Figure 1D, 1E), while amounts of pulled-down Arp1 and p50-S are not significantly changed (Figure 1D, 1E). In the Δ*vezA* background, *gpdA*-p50-S further decreases the amount of p150 pulled down by Arp11-GFP but the change in the amount of Arp1 or p50-S pulled down with Arp11-GFP is not statistically significant (Figure 1D, 1E).

The effects of p50 overexpression have been first studied in mammalian cells including neurons, and the results suggest that excess p50 can separate the shoulder from the mini-filament (Echeverri et al., 1996; Eckley et al., 1999; Karki et al., 1998; LaMonte et al., 2002; Maier et al., 2008; Melkonian et al., 2007). In vitro, a ∼25-fold excess of p50 leads to the release of two p150 molecules, two p50 molecules, and one p24 from the dynactin complex (Melkonian et al., 2007). However, a reduction in overall p150 protein levels has not been found in these systems. In *A. nidulans*, the pronounced decrease in p150 levels upon p50 overexpression may result from dissociation of p150 from the Arp1 mini-filament, as loss of Arp1 is known to reduce p150 levels in filamentous fungi and *Drosophila* (Haghnia et al., 2007; Minke et al., 1999; Zhang et al., 2024; Zhang et al., 2008).

It is known that a dynein or dynactin loss-of-function mutation would affect colony morphology in *A. nidulans*. However, strains containing the *gpdA*-p50-S allele form normal colonies, which is in contrast to the compact colonies formed by dynactin conditional-null mutants whose nuclear-distribution (nud) defect contributes to the colony defects (Figure 1F) (Xiang, 2018; Zhang et al., 2008). However, the *gpdA*-p50-S allele affects dynein-mediated early endosome distribution, as evidenced by an abnormal accumulation of mCherry-RabA-labelled early endosomes (Abenza et al., 2009; Zhang et al., 2010) near the hyphal tip in >50% of *gpdA*-p50-S hyphal-tip cells (Figure 1G).

In filamentous fungi, GFP-labelled dynein and dynactin form comet-like structures near the hyphal tip, representing microtubule-plus end accumulation (Figure 1H, 1I) (Han et al., 2001; Lenz et al., 2006; Xiang et al., 2000; Zhang et al., 2003). This plus-end accumulation of dynein or dynactin is conserved in budding yeast and in mammalian cells (Lee et al., 2003; Moore et al., 2008; Moughamian et al., 2013; Sheeman et al., 2003; Splinter et al., 2012; Valetti et al., 1999; Vaughan et al., 1999). In cells with *gpdA*-p50-S, dynactin p150-GFP comets are still present (Figure 1H), and the lowered intensity is expected as the p150 protein level is significantly decreased in the *gpdA*-p50-S background (Figure 1B, 1C). GFP-dynein comets are also present in the *gpdA*-p50-S background but the intensity is lower (Figure 1I, 1J). This is possibly resulted from the decrease in the p150 level since p150, especially its microtubule-binding domain, is critical for the plus-end accumulation of dynein (Duellberg et al., 2014; Yao et al., 2012; Zhang et al., 2003).

### Overexpression of p50 in *A. nidulans* causes the dynactin pointed-end proteins to form clusters

Unexpectedly, the localization pattern of Arp11-GFP changed dramatically upon p50 overexpression (Figure 2A). In wild-type background, Arp11-GFP, like all dynactin components, forms comet-like structures near the hyphal tip, representing microtubule plus-end accumulation (Figure 2A). However, in the majority of the *gpdA*-p50-S cells, a bright cluster can be seen (Figure 2A). In ∼80% of the cells showing the clusters, a single cluster can be seen within 5 μm from the hyphal tip, but the cluster does not always co-localize with mCherry-RabA-marked early endosomes near the hyphal tip (Figure 2A). In ∼20% of cells, the cluster can be more than 5 μm away from the hyphal tip (Figure 2A), and some cells also showed two or more small clusters near hyphal tip. Cluster formation is not fully correlated with an abnormal accumulation of early endosomes near the hyphal tip. In one experiment with 54 randomly selected hyphal-tip cells, 50 of them showed clusters but only 40 of them showed both the clusters and the hyphal-tip accumulation of early endosomes. In a control strain containing p50-S under the control of the endogenous promoter, Arp11-GFP still forms plus-end comets (Figure 2B). Thus, the Arp11-GFP cluster formation in the *gpdA*-p50-S background is not due to the presence of the S-tag in p50. The intensity of the Arp11-GFP cluster in the *gpdA*-p50-S background is much higher than that of Arp11-GFP comets in the p50-S background (Figure 2C). Moreover, other GFP-labeled pointed-end proteins such a p62-GFP and p25-GFP proteins also form clusters in the *gpdA*-p50-S background (Figure 2D).

**Figure 2.**
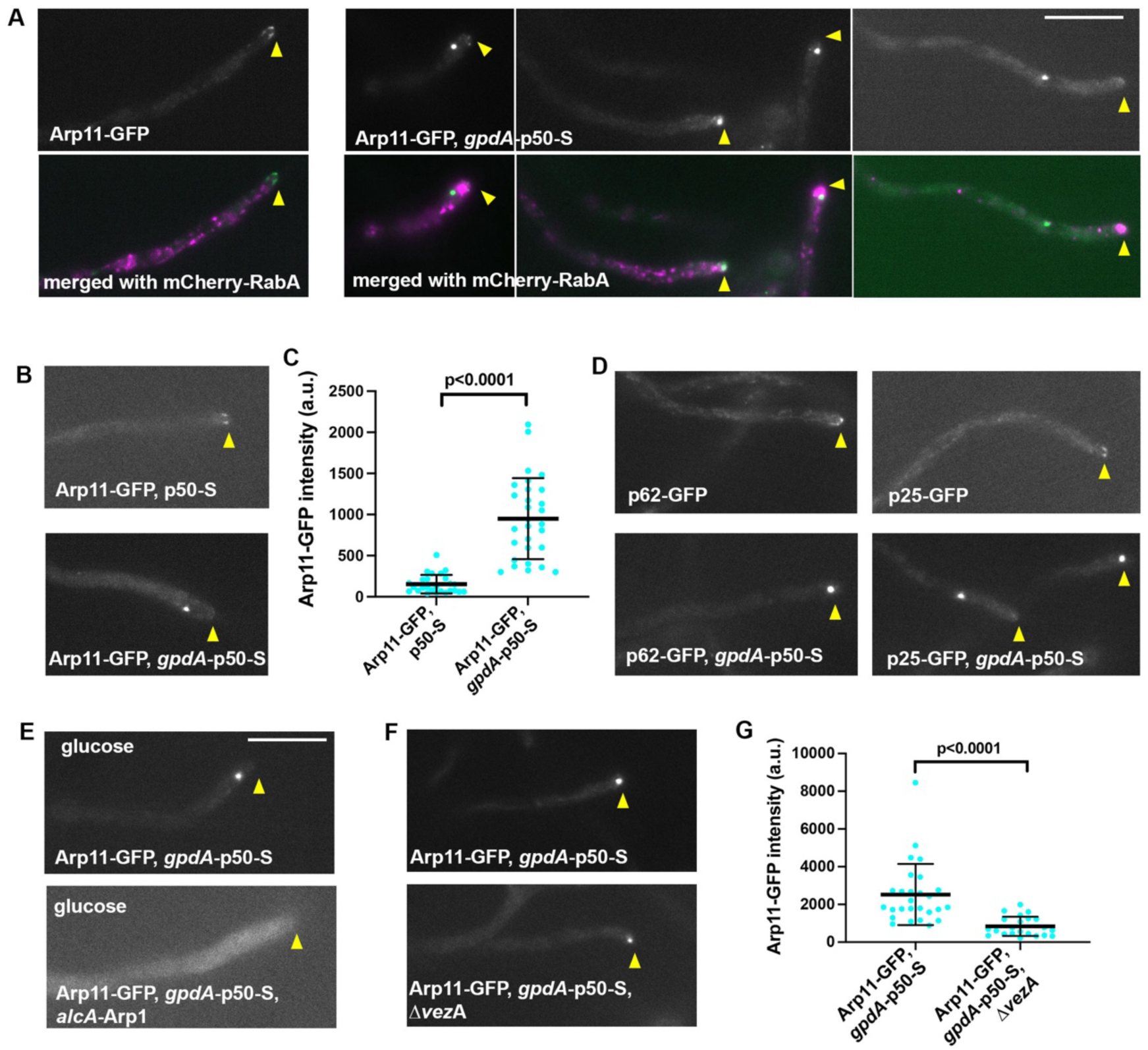
Dynactin pointed-end proteins form clusters in cells with p50 overexpression (*gpdA*-p50-S). (A) Arp11-GFP normally form plus-end comets but form bright clusters near the hyphal tip in the *gpdA*-p50-S background. Merged images showing GFP and mCherry-RabA-labeled early endosomes are presented below. Hyphal tip is indicated by a yellow arrowhead. Bar, 10 μm. (B) Arp11-GFP in p50-S and *gpdA*-p50-S backgrounds. (C) A quantitative analysis of Arp11-GFP signals in p50-S and *gpdA*-p50-S backgrounds. Scatter plots with mean and S.D. values were generated by Prism 10, and the p values were generated by Mann-Whitney test (unpaired). (D) Images of p62-GFP and p25-GFP in wild-type and *gpdA*-p50-S backgrounds. (E) The Arp11-GFP cluster formed in the *gpdA*-p50-S cells could not be observed in the *alcA*-Arp1 background where the expression of Arp1 is shut off on glucose. (F) Arp11-GFP clusters in the *gpdA*-p50-S and *gpdA*-p50-S, Δ*vezA* cells. (G) A quantitative analysis of Arp11-GFP cluster intensity. Scatter plots with mean and S.D. values were generated by Prism 10, and the p values were generated by Mann-Whitney test (unpaired).

We next determined if cluster formation by the pointed-end proteins upon p50 overexpression depends on Arp1 by using *alcA*-Arp1, a conditional Arp1-null allele (Zhang et al., 2024) (note that the *alcA*-promoter-driven Arp1 expression is shut off on glucose) (Waring et al., 1989). In the strain containing both *gpdA*-p50-S and *alcA*-Arp1, Arp11-GFP signals are diffuse in glucose medium (Figure 2E). Thus, Arp1 is essential for Arp11-GFP cluster formation upon p50 overexpression.

We further determined if VezA plays any role in Arp11-GFP cluster formation upon p50 overexpression. In the strain containing both *gpdA*-p50-S and Δ*vezA*, Arp11-GFP still forms clusters (Figure 2F). However, the fluorescent intensity of the cluster is significantly reduced (Figure 2G), suggesting that VezA enhances Arp11 cluster formation.

### Overexpression of p50 in *A. nidulans* causes VezA-GFP to form clusters

We next examined VezA-GFP upon p50 overexpression. Astonishingly, *gpdA*-p50-S also causes VezA-GFP to form clusters similar to those formed by dynactin pointed-end proteins (Figure 3A). Thus, p50 overexpression significantly changes VezA-GFP localization, as VezA-GFP normally is enriched at the hyphal tip but does not form this type of clusters (Yao et al., 2015; Zhang et al., 2024). In most cases (∼80%), a small cluster can be seen within 5 μm from the hyphal tip, but a cluster far away from the hyphal tip could also be observed. VezA-GFP missing the transmembrane domain (ΔTM-VezA-GFP) also forms clusters but their intensity is lower than that of the full-length VezA-GFP (Figure 3A, 3B). Previously, we have constructed VezA^Δ1-20^-GFP (or ΔN-VezA-GFP) and VezA^Δ563-615^-GFP (or ΔC-VezA-GFP) alleles and showed that these mutants are defective in dynein-mediated early endosome transport (Zhang et al., 2024). This was consistent with the AlphaFold-based structural prediction that the N- and C-termini of VezA are important for interaction with the dynactin pointed end and with Arp1, respectively (Zhang et al., 2024). Here we found that ΔN-VezA-GFP or ΔC-VezA-GFP proteins are unable to form clusters in *gpdA*-p50-S cells (Figure 3C). These results are consistent with the hypotheses that the VezA-GFP cluster formation depends on the interaction of VezA with dynactin components, and that the N- and C-termini of VezA are important for this interaction. To further test the requirement for VezA-GFP cluster formation, we introduced the Arp1- or Arp11-conditional null alleles, *alcA*-Arp1 or *alcA*-Arp11, to strains containing VezA-GFP and *gpdA*-p50-S. We found that VezA-GFP could no longer form clusters upon p50 overexpression in the absence of Arp1 or Arp11 expression (Figure 3D).

**Figure 3.**
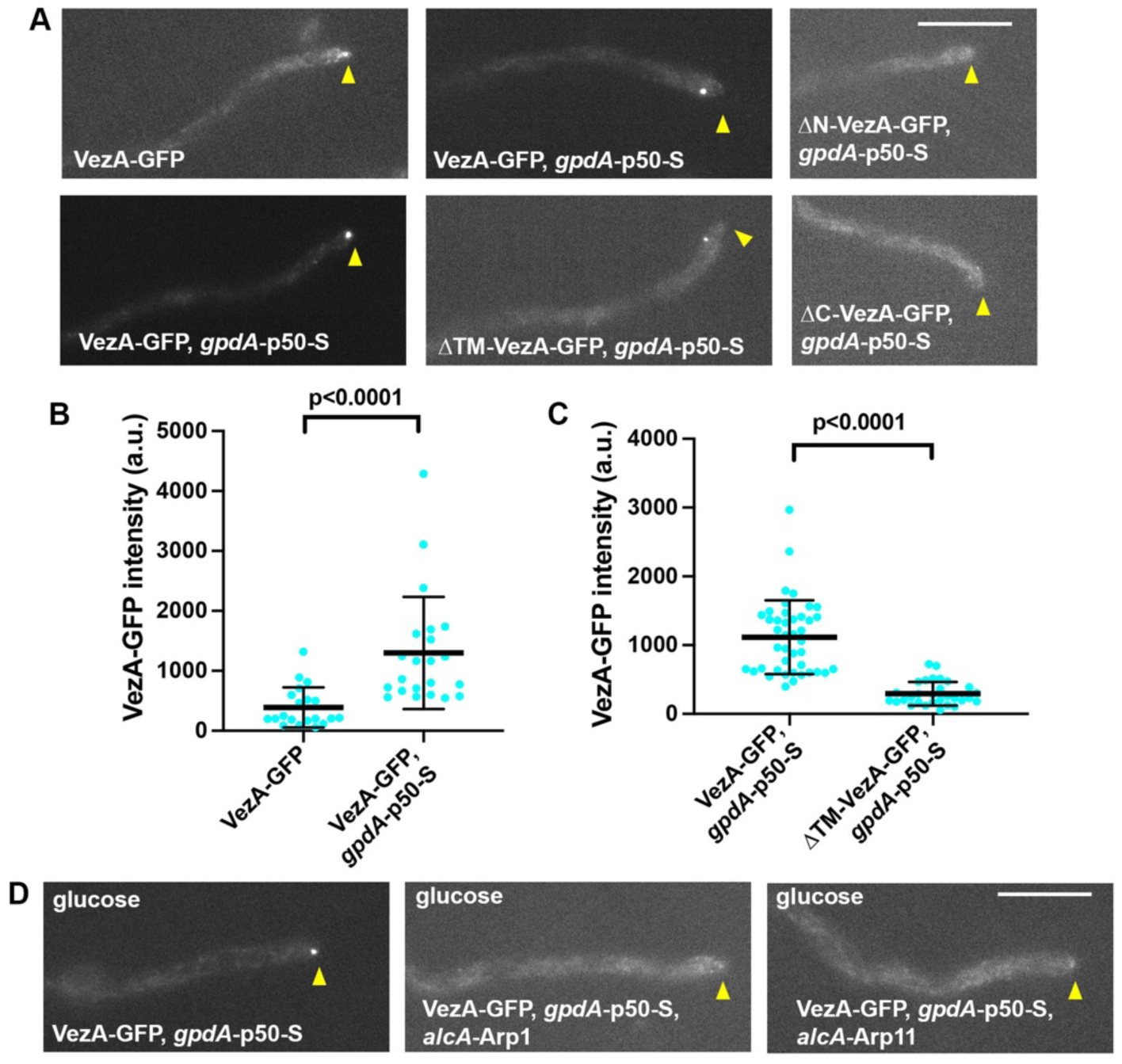
VezA-GFP forms clusters in p50-overexpressed cells. (A) While VezA-GFP signals are normally enriched at the hyphal tip, VezA-GFP forms clusters in the *gpdA*-p50-S background. ΔTM-VezA-GFP forms fainter clusters in the *gpdA*-p50-S background, but ΔN-VezA-GFP (VezA^Δ1-20^-GFP) or ΔC-VezA-GFP (VezA^Δ563-615^-GFP) does not form obvious clusters in the *gpdA*-p50-S background. Hyphal tip is indicated by a yellow arrowhead. Bar, 10 μm. (B) A quantitative analysis of VezA-GFP in wild-type and *gpdA*-p50-S backgrounds. Scatter plots with mean and S.D. values were generated by Prism 10, and the p values were generated by Mann-Whitney test (unpaired). (C) A quantitative analysis of VezA-GFP and ΔTM-VezA-GFP cluster intensity in the *gpdA*-p50-S background. Scatter plots with mean and S.D. values were generated by Prism 10, and the p values were generated by Mann-Whitney test (unpaired). (D) VezA-GFP can no longer form any cluster in the *gpdA*-p50-S background if the *alcA* promoter expressing Arp1 (*alcA*-Arp1) or Arp11 (*alcA*-Arp11) is shut off by glucose. Hyphal tip is indicated by a yellow arrowhead. Bar, 10 μm.

### Overexpressed p50-GFP form clusters in an Arp1-dependent fashion

Next, we constructed a strain containing *gpdA*-p50-GFP to observe overexpressed p50-GFP itself. We found that *gpdA*-p50-GFP is no longer localized at comet-like structure (representing microtubule plus-end accumulation) but forms clusters near the hyphal tip on top of a diffuse background of cytoplasmic fluorescence (Figure 4A). Importantly, shutting off the expression of Arp1 in the strain containing *alcA*-Arp1 grown on glucose eliminated cluster formation (Figure 4A), while shutting off Arp11 expression (*alcA*-Arp11 on glucose) increased the fluorescent intensity of the p50-GFP clusters (Figure 4A, 4B).

**Figure 4.**
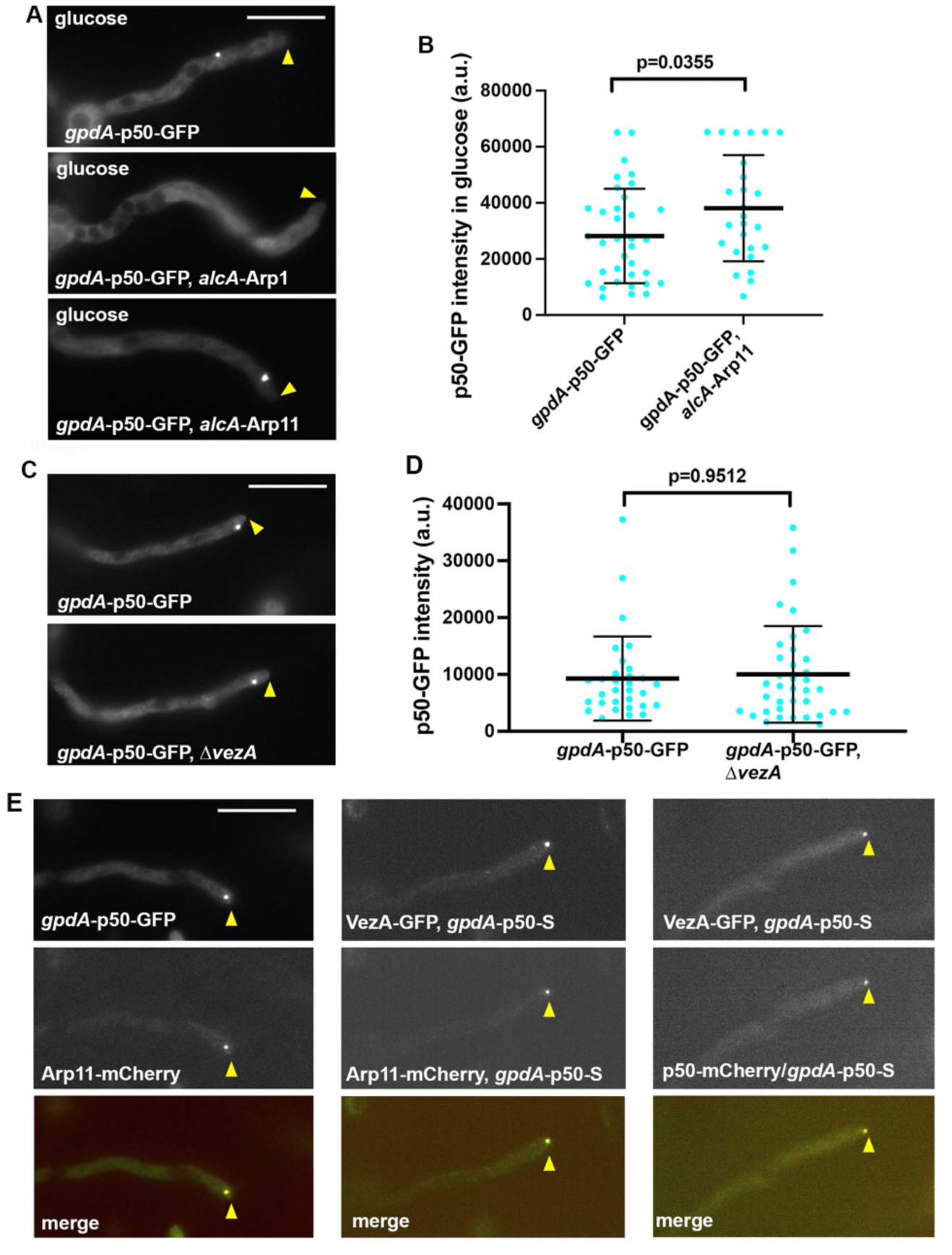
Overexpressed p50-GFP (*gpdA*-p50-GFP) form clusters in an Arp1-dependent and VezA-independent fashion, and the p50, Arp11 and VezA clusters are co-localized. (A) Images of *gpdA*-p50-GFP clusters in glucose medium. No clusters are observed in *alcA*-Arp1 cells. However, cells with *alcA*-Arp11 show bigger clusters. Hyphal tip is indicated by a yellow arrowhead. Bar, 10 μm. (B) A quantitative analysis of *gpd*A-p50-GFP and *gpd*A-p50-GFP, alcA-Arp11 cluster intensity. Scatter plots with mean and S.D. values were generated by Prism 10, and the p values were generated by Welch t-test (unpaired). (C) The p50-GFP cluster formation in p50-overexpressed cells with or without VezA. (D) A quantitative analysis of p50-GFP cluster intensity in *gpdA*-p50-S-containing cells with or without VezA (Δ*vezA*). Scatter plots with mean and S.D. values were generated by Prism 10, and the p values were generated by Mann-Whitney test (unpaired). (E) VezA, Arp11 and p50 are co-localized in p50-overexpressed cells. The left panel shows that in a cell with the *gpdA*-p50-S allele, Arp11-mCherry forms a cluster that co-localizes with the VezA-GFP cluster. The middle panel shows that in a cell with the *gpdA*-p50-GFP allele, Arp11-mCherry forms a cluster that co-localizes with that formed by p50-GFP. The right panel shows co-localization of a VezA-GFP cluster with a p50-mCherry cluster in a diploid containing both the *gpdA*-p50-S and p50-mCherry alleles in the p50 gene locus. Hyphal tip is indicated by a yellow arrowhead. Bar, 10 μm.

We next introduced the Δ*vezA* allele into the *gpdA*-p50-GFP strain to determine if VezA plays any role in p50-GFP cluster formation. However, we found that VezA does not significantly change the intensity of the p50-GFP cluster (Figure 4C, 4D). This is in contrast to VezA being important in enhancing Arp11-GFP cluster formation upon p50 overexpression (2F, 2G).

### VezA colocalizes with both p50 and Arp11 upon p50 overexpression

We next sought to determine if p50-GFP, Arp11-GFP, and VezA-GFP are in the same clusters upon p50 overexpression. To do that, we first generated a strain containing Arp11-mCherry and introduced *gpdA*-p50-GFP or *gpdA*-p50-S into this strain. Arp11-mCherry forms clusters upon overexpression of p50-GFP or p50-S (Figure 4E; Figure S1). In strains with both *gpdA*-p50-GFP and Arp11-mCherry, the p50-GFP and Arp11-mCherry signals were co-localized (Figure 4E). More importantly, we constructed a strain with VezA-GFP, Arp11-mCherry and *gpdA*-p50-S, and we found that the VezA-GFP cluster co-localizes perfectly with the Arp11-mCherry cluster (Figure 4E). In addition, we also determined if VezA co-localizes with p50 clusters formed upon overexpression. To do this, we first created a p50-mCherry strain and combined the p50-mCherry with the *gpdA*-p50-S and VezA-GFP alleles by making a diploid strain. In this diploid, p50-S overexpression causes both endogenously expressed p50-mCherry and VezA-GFP to form clusters and the signals were also co-localized (Figure 4E). These results indicate that these proteins are in the same clusters formed upon p50 overexpression.

### Loss of p150 or loss of an Arp1-binding region of p50 does not cause Arp11-GFP or VezA-GFP to form clusters

Since p50 overexpression causes a decrease in p150 level, which is likely because of its separation from the complex (Cheong et al., 2014; Echeverri et al., 1996; LaMonte et al., 2002; Melkonian et al., 2007), we examined if removing p150 will lead to Arp11-GFP cluster formation. We found that Arp11-GFP does not form clusters if p150 expression is repressed in the *alcA*-p150 background; the signals are diffuse, although the diffuse signals are slightly brighter at the hyphal tip region (Figure S2). Thus, cluster formation does not simply happen in response to a reduction in p150 protein level.

We next sought to test if any abnormality in the p50-Arp1 interaction would lead to the formation of the abnormal clusters containing Arp11 and VezA. The N-termini of p50 form extended regions (or “tentacles”) that interact with Arp1 subunits (Cheong et al., 2014; Urnavicius et al., 2015). In the cryo-EM structure of porcine dynactin (PDB 8PTK), aa19-21 from four different p50 subunits interact with four different Arp1 subunits via beta-sheet structures, but aa47-49 and aa87-89 also form beta-sheet structures with three other Arp1s (Figure 5A, 5B; Figure S3) (Singh et al., 2024; Urnavicius et al., 2015). The first Arp1-binding site is highly conserved, and Alphafold-3-based analysis identified this binding site in *A. nidulans* p50 (Figure 5B, 5C; Figure S3). To test the consequence of removing this site, we made the p50^Δ17-25^-S strain. This strain shows an obvious colony defect similar to that of the conditional-null mutant of p50, *alcA*-p50 (Figure 5D), indicating that this site is critical for p50 function. The p50^Δ17-25^-S mutant also shows a defect in dynein-mediated early endosome transport (Figure 5E), but it shows no clusters of Arp11-GFP or VezA-GFP (Figure 5F).

**Figure 5.**
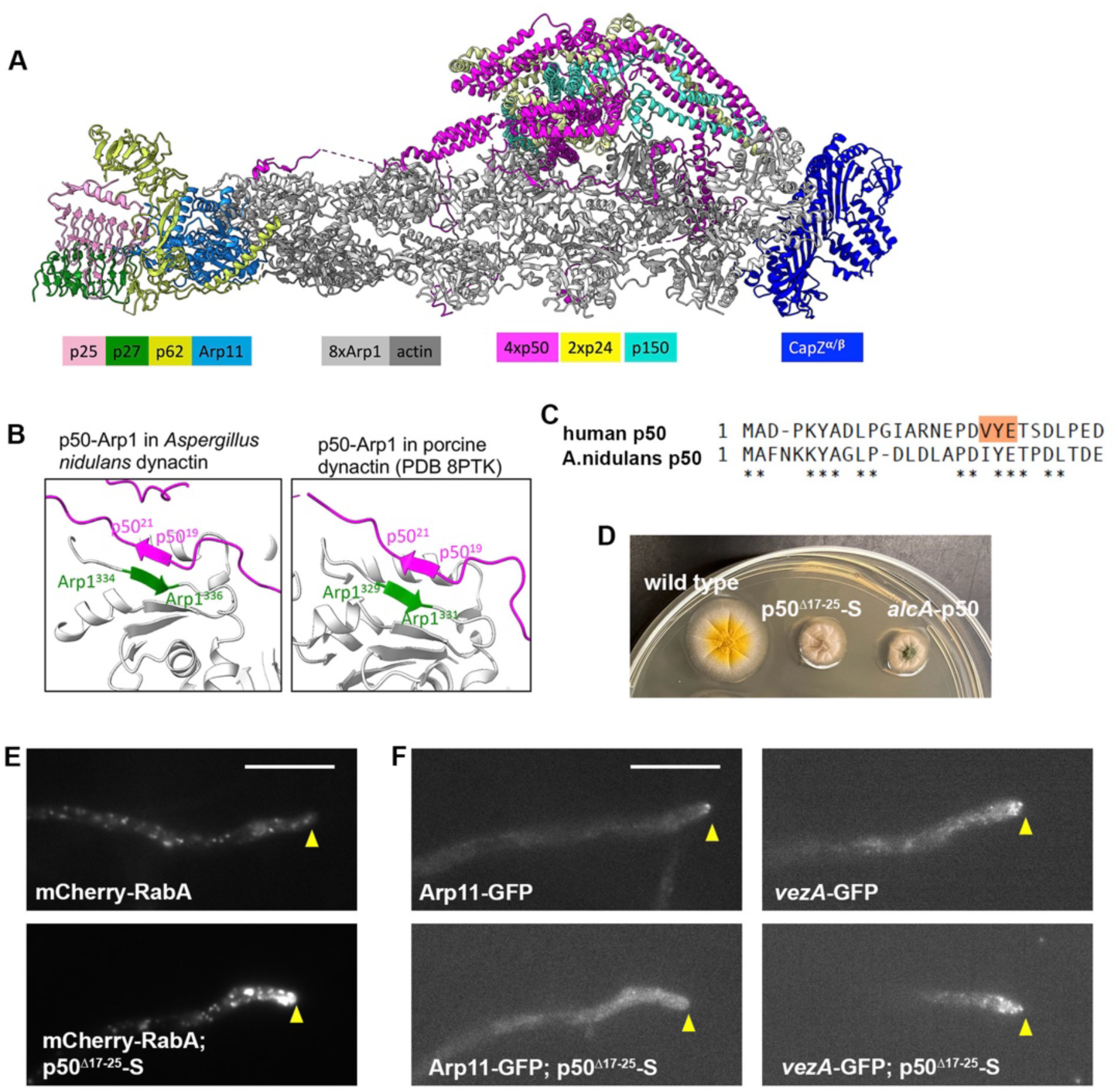
Phenotype of the p50^Δ17-25-^S mutant lacking an Arp1-binding site. (A) Cryo-EM structure of mammalian dynactin (source: PDB:8PTK (Singh et al., 2024)). For p150, only the C-terminus is shown. (B) A beta-sheet structure formed by p50-Arp1. The *A. nidulans* p50-Arp1 interaction is predicted by Alphafold3. The p50-Arp1 interaction in porcine dynactin is based on the published structure PDB:8PTK (Singh et al., 2024)). Note that Arp1^333^ in *A. nidulans* Arp1 corresponds to Arp1^328^ in human Arp1. (C) A small portion of the sequence alignment showing the N-termini of human p50 and *A. nidulans* p50. Note that the amino acids (highlighted in human p50) involved in the interaction with Arp1 are largely conserved (please see Figure S3 for the whole alignment). (D) The p50^Δ17-25-^S strain shows a colony defect similar to that of *alcA*-p50, a strain in which p50 expression is repressed by glucose. (E) In the p50^Δ17-25-^S mutant, mCherry-RabA-labeled early endosomes accumulate abnormally at the hyphal tip region. Hyphal tip is indicated by a yellow arrowhead. Bar, 10 μm. (F) Arp11-GFP and VezA-GFP localizations with or without the p50^Δ17-25-^S mutation. Note that neither Arp11-GFP nor VezA-GFP forms clusters in the p50^Δ17-25-^S mutant background. Hyphal tip is indicated by a yellow arrowhead. Bar, 10 μm.

### Loss of aa17-25 of p50 eliminates the Arp11, VezA and p50 clusters upon p50 overexpression

Given that Arp1 is critical for the formation of p50-GFP, Arp11-GFP and VezA-GFP clusters upon p50 overexpression, we believe that Arp1 is a core component of the cluster (although we have not been able to construct a GFP-labeled Arp1 fusion that is functional). We next sought to test whether the p50-Arp1 interaction within the cluster is mediated by aa17-25 of p50 involved in the Arp1-p50 beta sheet structures (Figure 5A, 5B; Figure S3) (Singh et al., 2024; Urnavicius et al., 2015). To do that, we created a *gpdA*-p50^Δ17-25^-S allele in the p50 gene locus to cause overexpression of the mutant p50, and we also introduced Arp11-GFP into this strain background. In contrast to Arp11-GFP clustering in the *gpdA*-p50-S background, there is no cluster of Arp11-GFP upon p50 overexpression if the conserved Arp1-binding site is removed (Figure 6A). The Arp11-GFP signals are diffuse in the *gpdA*-p50^Δ17-25^-S background although the diffuse signals are slightly brighter at the hyphal tip region (Figure 6A). Similarly, we also introduced VezA-GFP into the *gpdA*-p50^Δ17-25^-S background and found no obvious clusters of VezA-GFP upon p50 overexpression if the conserved Arp1-binding site is removed (Figure 6B).

**Figure 6.**
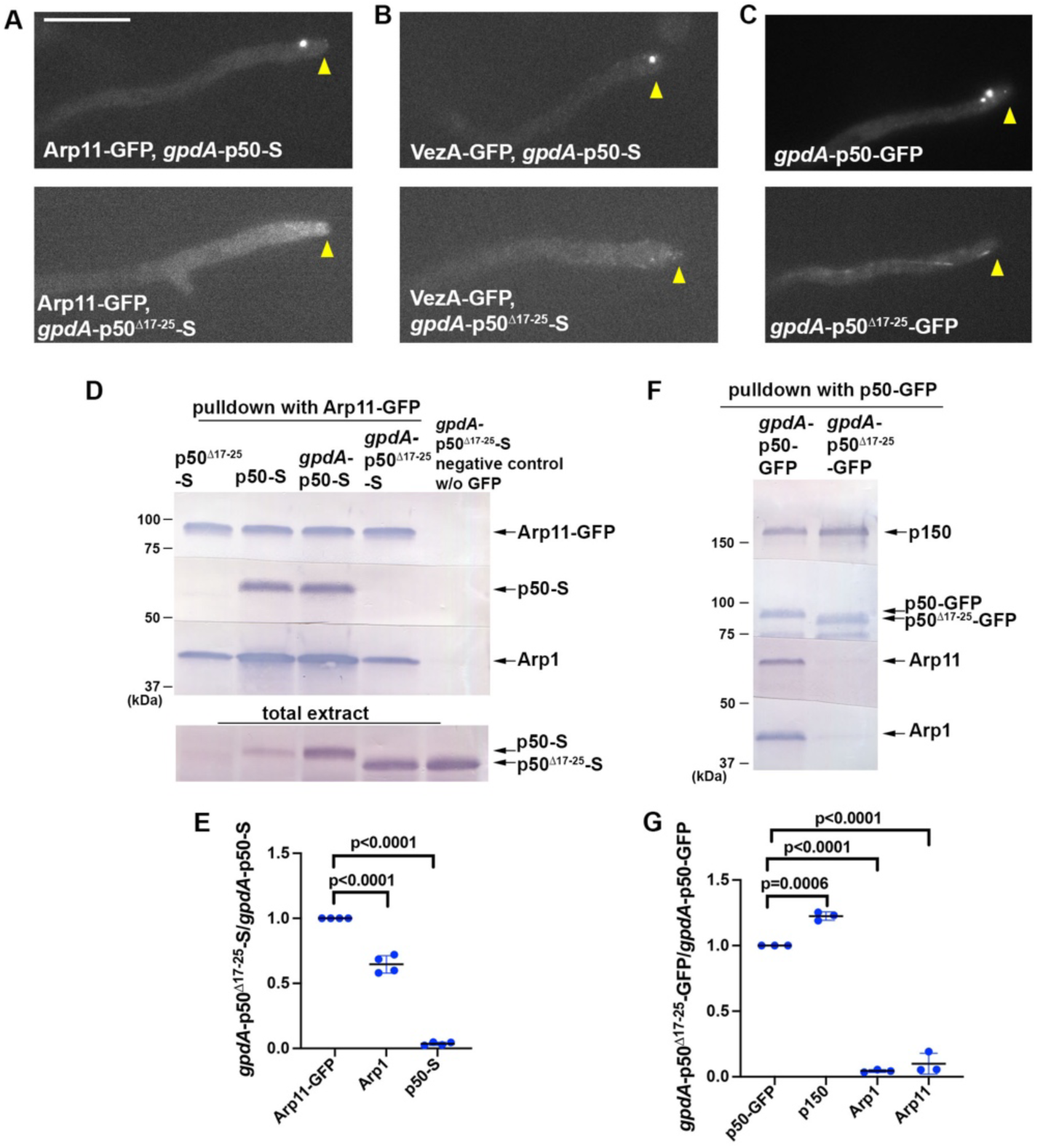
The p50 amino acids 17-25 is required for the formation of Arp11-GFP, VezA-GFP and p50-GFP clusters upon p50 overexpression. (A) Arp11-GFP forms clusters in the *gpdA*-p50-S background but not in the *gpdA*-p50^Δ17-25^-S background. Hyphal tip is indicated by a yellow arrowhead. Bar, 10 mm. (B) VezA-GFP forms clusters in the *gpdA*-p50-S background but not in the *gpdA*-p50^Δ17-25^-S background. Hyphal tip is indicated by a yellow arrowhead. (C) p50-GFP forms clusters in the *gpdA*-p50-GFP background but not in the *gpdA*-p50^Δ17-25^-GFP background. Note that signals in the *gpdA*-p50^Δ17-25^-GFP background appear to be enriched at microtubule plus ends. Hyphal tip is indicated by a yellow arrowhead. (D) Western blots showing that p50-S could hardly be pulled down with Arp11-GFP in the *gpdA*-p50^Δ17-25^-S strain. The p50-S signals from the total extracts used for the same pulldown experiment are shown at the bottom. The strains used for the experiment all contain Arp11-GFP except for the negative control. (E) A quantitative analysis on the effect of *gpdA*-p50^Δ17-25^-S on the amounts of Arp1 and p50-S pulled down with Arp11-GFP. The ratios of the intensity values of *gpdA*-p50^Δ17-25^-S to those of *gpdA*-p50-S are shown for each component, and these ratios are relative to the ratio for the Arp11-GFP that was set as 1. Scatter plots with mean and SD values as well as p values were generated by Prism 10 (Ordinary one-way ANOVA with Dunnett’s multiple comparison test). (F) Western blots showing that overexpressed p50^Δ17-25^-GFP (*gpdA*-p50^Δ17-25^-GFP) pulls down p150 but much lower amounts of Arp1 and Arp11 compared to those pulled down with overexpressed p50-GFP. (G) A quantitative analysis on the effect of *gpdA*-p50^Δ17-25^-GFP on the amounts of p150, Arp1 and Arp11 pulled down with p50-GFP. The ratios of the intensity values of *gpdA*-p50^Δ17-25^-GFP to those of *gpdA*-p50-GFP are shown for each component, and these ratios are relative to the ratio for p50-GFP that was set as 1. Scatter plots with mean and SD values as well as p values were generated by Prism 10 (Ordinary one-way ANOVA with Dunnett’s multiple comparison test).

To observe overexpressed p50-GFP directly in a strain without the Arp1-binding site, we created an allele of *gpdA*-p50^Δ17-25^-GFP in the p50 gene locus. In contrast to what we have observed in the *gpdA*-p50-GFP background, there is no cluster of p50-GFP in the *gpdA*-p50^Δ17-25^-GFP strain (Figure 6C). However, faint microtubule decoration can be seen (Figure 6C), similar to the localization of p150 upon p50 overexpression (See Figure 1H). This result suggests that overexpressed p50^Δ17-25^-GFP associates with p150, and their microtubule binding is via the microtubule-binding domain of p150 (Waterman-Storer et al., 1995).

To further examine the consequence of deleting aa17-25 of p50 in the context of p50 overexpression, we performed biochemical pulldown assays. We first determined if p50-S can be pulled down with Arp11-GFP in the p50^Δ17-25^-S (endogenous p50 promoter) or the *gpdA*-p50^Δ17-25^-S background. In the p50^Δ17-25^-S strain, the protein level of p50^Δ17-25^-S in total extract is obviously lower than that of p50-S in the p50-S strain. In the *gpdA*-p50^Δ17-25^-S strain where the mutant p50 is overexpressed, the level of p50^Δ17-25^-S in total cell extract is similar to that of p50-S in the *gpdA*-p50-S-containing strain (Figure 6D), but we could barely detect any p50^Δ17-25^-S pulled down with Arp11-GFP (Figure 6D, 6E). In addition, while Arp1 can be pulled down, the amount of pulled-down Arp1 is significantly reduced in the mutant (Figure 6D, 6E), similar to the effect caused by shutting off p50 expression (Zhang et al., 2024). We next performed pulldown experiments using overexpressed p50-GFP and p50^Δ17-25^-GFP. While the amount of p150 pulled down with overexpressed p50^Δ17-25^-GFP is moderately increased, the amount of Arp11 or Arp1 pulled down with overexpressed p50^Δ17-25^-GFP is significantly decreased compared to that in the *gpdA*-p50-GFP control (Figure 6F, 6G). Together, these data provide strong evidence for the importance of this small region of p50 in dynactin assembly.

## Discussion

The dynactin complex is made of the Arp1 mini-filament capped by capping protein and the pointed-end subcomplex containing Arp11, p62, p25 and p27, and the shoulder sub-complex containing p150, p50 and p24 (Schroer, 2004). While studies have revealed the structure of the mature complex, the assembly process still needs investigation (Chaaban and Carter, 2022; Chowdhury et al., 2015; Imai et al., 2014; Lau et al., 2021; Schafer et al., 1994; Singh et al., 2024; Urnavicius et al., 2015). It has been known that excess p50 separates the dynactin shoulder from the mini-filament (Echeverri et al., 1996; Eckley et al., 1999; Karki et al., 1998; LaMonte et al., 2002; Maier et al., 2008; Melkonian et al., 2007). Here we found that excess p50 induces formation of novel Arp1-dependent clusters containing p50 and pointed-end proteins including Arp11, p62 and p25. Astonishingly, a recently identified dynactin assembly factor VezA (Zhang et al., 2024), which normally does not co-localize with dynactin in live cells, is present in these clusters upon p50 overexpression.

The p50 protein has an intrinsic tendency to self-assemble, but its binding to a smaller shoulder protein p24 prevents oligomerization in vitro (Maier et al., 2008). Thus, it is possible that p50 overexpression increases the p50-to-p24 ratio, thereby enhancing p50 oligomerization. In addition, as p24 is known to be critical for the p50-p150 interaction (Amaro et al., 2008; Terasawa et al., 2010), a decreased p24-to-p50 ratio should weaken the p150-p50 association and leave the p150-binding surface of p50 unoccupied, which may further enhance p50 oligomerization. As cluster formation from both p50-GFP and Arp11-GFP is strictly Arp1-dependent, Arp1 is likely another core component of the clusters. In this study, we have also found that a small region of p50 (aa17-25), which is involved in binding Arp1 proteins via beta-sheet structures (Singh et al., 2024; Urnavicius et al., 2015), is critical for p50-overexpression-driven formation of p50, Arp11 and VezA clusters.

Arp11 is not needed for the p50-overexpression-driven p50-GFP cluster formation, and unexpectedly, shutting off the expression of Arp11 significantly enhances the fluorescence intensity of p50-GFP clusters. It has been found previously that loss of Arp11 expression has a significant effect on Arp1 mini-filament assembly in *A. nidulans* and mammalian cells (Yeh et al., 2012; Zhang et al., 2024; Zhang et al., 2008). In the most recent experiments in *A. nidulans*, loss of Arp11 causes the amount of Arp1 pulled down with p50-GFP to be dramatically decreased to <25% of the normal level (Zhang et al., 2024). Thus, the pointed-end sub-complex may initiate Arp1 mini-filament assembly (Zhang et al., 2024). However, the cluster formation in the absence of Arp11 suggests an alternative p50-driven Arp1 assembly mechanism, similar to what has been proposed previously (Urnavicius et al., 2015). While the p50 protein is unstable upon loss of Arp1 expression, p50 remains stable upon loss of Arp11 expression, although loss of Arp11 expression causes a significant decrease in the amount of p50-associated Arp1 (Zhang et al., 2024). Although we still do not understand why loss of Arp11 makes the p50-GFP cluster brighter upon p50 overexpression, we speculate that the p50-Arp1 assembly in the absence of the pointed-end sub-complex is abnormal and may be more likely to aggregate with each other.

While the N-terminal extended region of p50 is known to be important for binding Arp1 directly (Cheong et al., 2014; Urnavicius et al., 2015), a functional analysis of the small regions involved in forming the p50-Arp1 beta-sheet structures has never been performed previously. We found that deleting aa17-25 of p50 causes a phenotype similar to that exhibited by a p50-conditional null mutant (*alcA*-p50). In the strain with overexpressed p50^Δ17-25^ protein, the protein level of mutant p50 is close to normal, and it interacts normally with p150, but its interaction with Arp1 is significantly reduced. Thus, although two other small regions of p50, aa47-49 and aa87-89, in mammalian dynactin also form beta-sheet structures with three different Arp1s (Singh et al., 2024; Urnavicius et al., 2015), the beta-sheet strand within aa17-25 plays a key role in holding the Arp1 mini-filament together. Moreover, Arp11-GFP can still pulldown Arp1 proteins in cells with p50^Δ17-25^-S, but the amount of Arp1 pulled down with Arp11-GFP is reduced, similar to what has been observed upon loss of p50 expression (Zhang et al., 2024). This result is consistent with the hypothesis that p50-Arp1 interactions ensure that the Arp1 mini-filament is assembled with the correct length (Urnavicius et al., 2015), but it also indicates that partial assembly of the Arp1 mini-filament linked to the pointed-end sub-complex can be achieved in the absence of the p50-Arp1 interactions.

The VezA cluster formation upon p50 overexpression requires Arp1, Arp11, the N-terminal 1-20 residues of VezA predicted to interact with the pointed-end sub-complex and the C-terminal 563-615 residues of VezA predicted to interact with Arp1. Thus, consistent with our previous biochemical and structural prediction data (Zhang et al., 2024), our current imaging data also places VezA at the interface between the Arp1 mini-filament and the pointed-end sub-complex. Moreover, we found that VezA significantly enhances the fluorescence intensity of Arp11-GFP clusters but not that of the p50-GFP clusters. These observations suggest that p50 and Arp1 initiate cluster formation upon p50 overexpression, followed by recruitment of Arp11-containing pointed-end sub-complex in a process partially facilitated by VezA. This partial dependence on VezA is consistent with VezA’s role as a non-essential but important dynactin assembly factor that strengthens the link between the pointed-end sub-complex and the mini-filament. However, given that the pointed-end sub-complex may initiate a partial assembly of the Arp1 mini-filament, it is also possible that VezA helps to connect this pointed-end sub-complex-linked mini-filament with p50-multimers via the p50-Arp1 interactions. Future studies will be needed to further test these possibilities. In normal cells, VezA must also be involved in dynactin shoulder assembly onto the mini-filament (Zhang et al., 2024): In the absence of VezA, the protein levels of p50 and p150 are reduced, but this reduction was not observed upon loss of Arp11. It is possible that in cells with p50 overexpression, this shoulder assembly function of VezA is not critical for cluster formation because the increased p50 concentration would facilitate the p50-Arp1 interactions. Although VezA was identified as a dynactin assembly factor, we have never observed its co-localization with dynactin. The fact that VezA-GFP are co-localized with Arp11 and p50 in the same clusters suggest that VezA is trapped in this assembly state driven by excess p50. How VezA normally dissociates from the dynactin complex will require further studies.

Combining all the results, we propose a model in which p50 oligomerization together with the normal p50-Arp1 interactions drive the formation of the p50-Arp1 assemblies (Figure 7). These assemblies subsequently recruit the pointed-end sub-complex containing Arp11, p62, p25 and p27, or the pointed-end-sub-complex linked to a few Arp1 subunits. Because Arp11-GFP pulls down normal amounts of Arp1 and p50 upon p50 overexpression (Figure 1D, 1E), we hypothesize that Arp1 proteins may still form mini-filaments in these clusters. VezA binds Arp1 and the pointed-end sub-complex. In the absence of VezA (Δ*vezA*), fewer pointed-end sub-complexes are recruited to the cluster, and any Arp1 mini-filament missing the pointed-end sub-complex may also miss some Arp1 subunits. Future studies will be needed to elucidate the detailed composition of the clusters, which may lead to a better understanding of the dynactin assembly process including the possible identification of other co-localized dynactin assembly factors.

**Figure 7.**
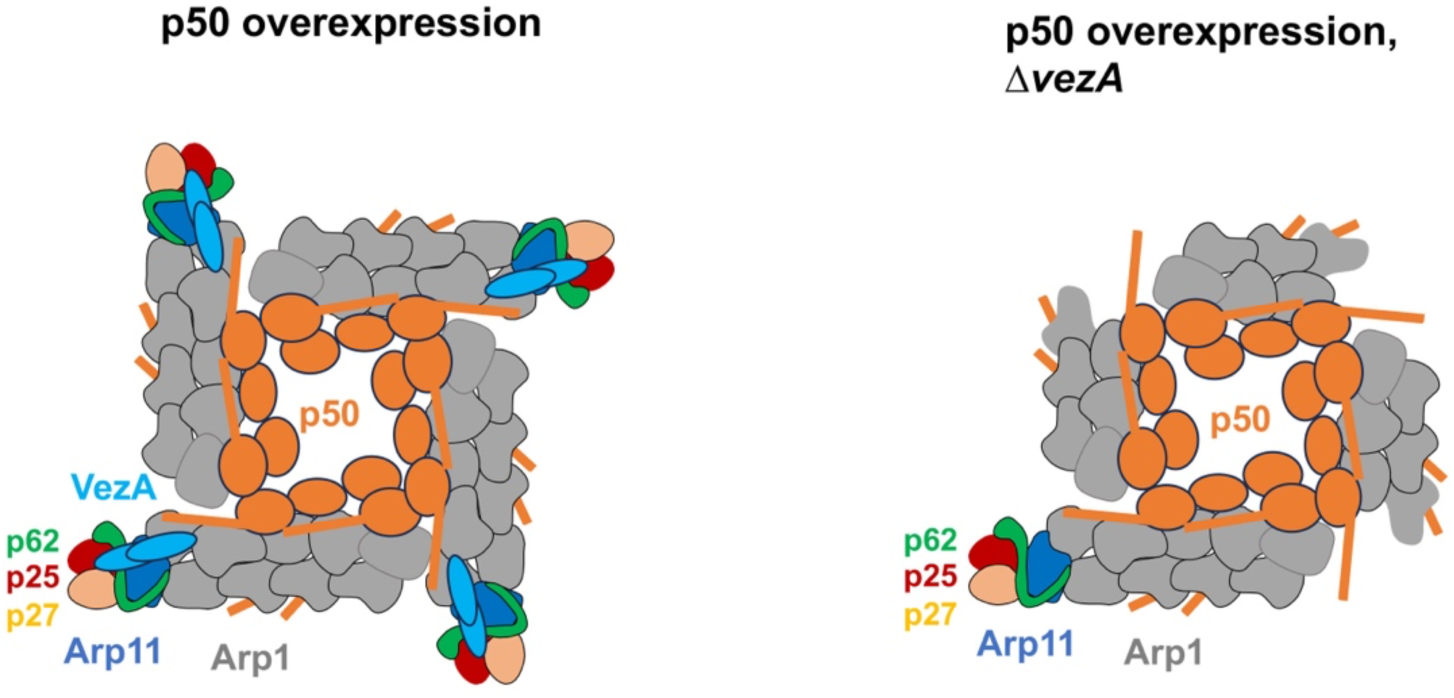
Models of p50-overexpression-driven formation of p50, Arp1 and VezA clusters in wild-type *vezA* background (left) and in the Δ*vezA* mutant background (right). In both situations, overexpressed p50 form oligomers, driving cluster formation. The elongated part of p50 represents the extended region (or the tentacle-like structure) formed by the N-terminal ∼100 amino acids, and these tentacles are responsible for interacting with Arp1 (Cheong et al., 2014; Urnavicius et al., 2015). For simplicity, the amino acids forming beta-sheet structures with Arp1 (Singh et al., 2024; Urnavicius et al., 2015) are not depicted in this cartoon. VezA is recruited to the cluster by Arp1 and the pointed-end sub-complex containing Arp11, p62, p25 and p27. In the Δ*vezA* background, there is a decrease in the number of pointed-end proteins in the cluster, and the integrity of those Arp1 mini-filaments missing the pointed-end sub-complex may also be compromised.

p50 overexpression in mammalian neurons causes late-onset neurodegeneration reminiscent of ALS (LaMonte et al., 2002). We note that the defect in dynein function caused by excess p50 is moderate in *A. nidulans*. Even though there is a clear defect in dynein-mediated early endosome transport, there is no obvious defect in colony formation, suggesting that dynein-mediated nuclear migration is normal (Xiang, 2018). In *A. nidulans*, early endosome transport is more sensitive than nuclear migration to dynein deficiency, because a low level of dynein activity can support nuclear migration but not early endosome transport (Tan et al., 2014). Thus, a p50-overexpression-driven decrease in the level of functional dynactin complexes that does not obviously affect nuclear movement or colony growth could affect early endosome transport. In this context, it should also be pointed out that dynein mutations identified from malformations of cortical development (involving nuclear-migration-participated neuronal migration (Morris et al., 1998; Tsai and Gleeson, 2005)) tend to cause more severe dynein defects in vitro or in model organisms compared to dynein mutations identified from neurodegenerative diseases (Hafezparast et al., 2003; Hoang et al., 2017; Marzo et al., 2019; Ori-McKenney et al., 2010; Salvador-Garcia et al., 2024; Sivagurunathan et al., 2012). In the case of p50 overexpression in mammalian neurons, reduction in functional dynactin availability could compromise long-distance transport, contributing to neurodegenerative phenotypes (LaMonte et al., 2002). On the other hand, formation of molecular clusters may also contribute to neuronal degeneration; for example, an ALS-causing kinesin-1 mutation resulting in motor activation also causes motor aggregation (Baron et al., 2022; Nakano et al., 2022; Pant et al., 2022). It should be interesting to determine whether similar clusters of dynactin components are formed in neurons or other cell types upon p50 overexpression.

## Materials and Methods

### *A. nidulans* strains, media and general methods

*A. nidulans* strains used in this study are listed in Table S1. *A. nidulans* strains grow on solid rich medium made of either YAG (0.5% yeast extract and 2% glucose with 2% agar) or YAG+UU (YAG plus 0.12% uridine and 0.11% uracil) (Qiu et al., 2023). Solid minimal medium containing 1% glucose was used for selecting progeny from a genetic cross. Note that the minimal medium (pH 6.5) also contains 0.6% NaNO_3_, 0.052% KCl, 0.0152% KH_2_PO_4_ and 0.051% MgSO_4_. For most live-cell imaging experiments, cells were cultured overnight at 32°C in a liquid minimal medium containing 1% glycerol. For the live-cell imaging experiments involving the *alcA*-promoter-based conditional mutants such as *alcA-*Arp1, *alcA*-Arp11, *alcA*-p150 and *alcA*-p50 conditional-null mutants, we harvested spores from the solid minimal medium containing 1% glycerol (non-repressive for the *alcA* promoter) and cultured them in minimal medium containing 0.1% glucose (repressive for the *alcA* promoter) overnight at room temperature followed by 32°C for 5 hours before imaging. For western analyses or pulldown experiments, liquid YG medium (same as YAG but without agar) was used, but minimal medium containing 1% glycerol or 0.1% glucose was used for some experiments using strains containing *alcA*-based conditional mutants.

Strains containing multiple mutant alleles were constructed by genetic crosses. Genetic crosses were done by standard methods, and progeny with desired genotypes were selected based on colony phenotype, imaging analysis, western analysis, diagnostic PCR, and/or sequencing of specific regions of the genomic DNA. For genomic DNA isolation, about 0.25 g of hyphal mass was harvested from overnight culture for each sample and pulverized with liquid nitrogen by using mortars and pestles. The DNeasy Plant Mini Kit (QIAGEN) was used for genomic DNA isolation. The QIAquick Gel Extraction Kit (QIAGEN) was used for purifying PCR products from agarose gels. Sequencing was done using the service of Quintarabio (https://www.quintarabio.com/service/ngs_services).

### Live-cell imaging and analyses

Microscopic images used in most figures were generated using a Nikon Ti2-E inverted microscope with Ti2-LAPP motorized TIRF module and a CFI apochromat TIRF 100 x 1.49 N.A. objective lens (oil). The microscope was controlled by NIS-Elements software using 488 nm and 561 nm lines of LUN-F laser engine and ORCA-Fusion BT cameras (Hamamatsu). All the images were taken with a 0.1-s exposure time (binning: 2×2). Image labeling was done using Adobe Photoshop. For all images, cells were grown at 32°C for ∼18 hours in the LabTek Chambered #1.0 borosilicate coverglass system (Nalge Nunc International, Rochester, NY). Images were taken at room temperature. For quantitation of signal intensity, a region of interest (ROI) was selected and maximal intensity within the ROI was measured. Then the ROI box was dragged outside of the cell to take the background value, which was then subtracted from the intensity value. For measuring the signal intensity of microtubule plus-end comets formed by GFP-labeled dynein or dynactin components, only the comet closest to hyphal tip was measured.

### Construction of a strain containing *gpdA*-p50-S

The *gpdA* promoter fragment (1.23 kb) was amplified via PCR using genomic DNA from the RQ247 strain as a template with primers GPDF (5’-GACTCGAGTACCATTTAATTCTATTTG-3’) and GPDR (5’- TGTGATGTCTGCTCAAGCG -3’). To generate the *gpdA*-p50 sequence, two 1 kb fragments (N and C) were amplified from GR5 genomic DNA using primer pairs P50NN (5’-AGGGAGGTTTGAACCATGG-3’) / gpdFp50NC (5’-CAAATAGAATTAAATGGTACTCGAGTCAGCTAGAATATTGAAGGATCTTAGTTGTC-3’), and p50CC (5’- AAGTGCTTCAGCGTCTGCTG -3’) /gpdRp50ATG (5’-CCCGCTTGAGCAGACATCACAATGGCTTTCAACAAAAAATATGCTGGTC-3’), respectively. These three fragments, the gpdA promoter and the N/C p50 segments, were subsequently fused by PCR using primers p50NN1 (5’-TCGGAGATGGTTCGATCCTG-3’) and p50CC1 (5’-TTCCTGACGCGGGTAGAAAG -3’) to generate a 3.3 kb *gpdA-*p50 fusion product.

Separately, a 4 kb p50-*S-AfpyrG* fusion fragment was amplified from the genomic DNA of strain SX1 (Zhang et al. 2024) using primers p50GNN1 (5’- CGAGGCGAAGGACACATCA -3’) and p50GCC1 (5’- CATCTTTAACAGCTGCCGCC -3’). The 3.3 kb *gpdA*-p50 fragment and the 4 kb p50-*S-AfpyrG* fragment were then co-transformed into the XY42 strain. Transformants exhibiting high S-p50 expression levels were identified via Western blot and further validated through PCR sequencing using primers gpdAC5 (5’-CAGTTCGAGCTTTCCCACTT-3’) and AfpyrGN3 (5’-CGAAGAGGGTGAAGAGCATT-3’). The p50 coding region was sequenced to make sure that there are no amino-acid-changing mutations introduced accidently during the construction.

### Construction of the p50^Δ17-25-^S mutant

The p50^Δ17-25-^S mutant was constructed using a multi-step PCR and co-transformation strategy. Initially, two overlapping fragments flanking the deletion site—a 1.1 kb N-terminal fragment and a 0.9 kb C-terminal fragment—were amplified from *GR5* genomic DNA. This was achieved using the primers P50NdNC (5’-GACGGTGGAGGCTTCGTCTGTAGCGAGATCCTGGGCGAGT-3’), P50NN(5’-AGGGAGGTTTGAACCATGG-3’), P50NdCN (5’- ACAGACGAAGCCTCCACCG-3’) and P50CC (P50CC (5’- AAGTGCTTCAGCGTCTGCTG -3’). These two fragments were subsequently joined via fusion PCR to generate a 2.0 kb fragment containing the targeted Arp1-binding site mutation. In parallel, a 4.2 kb S-p50-ApyrG fusion fragment was amplified from the genomic DNA of the *SX1* strain using oligos p50GNN (5’- AAGATGAGATGGCGGCGTC -3’) and P50GCC (5’- TCCAACCACACAAGGAATG -3’) (Zhang et al., 2024). The resulting 2.0 kb and 4.2 kb PCR products were co-transformed into the XY42 strain. Transformants were screened and the p50 Arp1-binding site deletion, along with the presence of the S-tag, was confirmed through PCR and sequencing using primers P50CC1 (5’- TTCCTGACGCGGGTAGAAAG -3’) and S-tag3 (5’-GCTGGCGTTCGAATTTAGC-3’).

### Construction of a *gpdA*-p50^Δ17-25^-S strain

The construction of the *gpdA*-p50^Δ17-25^-S strain involved the co-transformation of two distinct PCR products into the RQ54 host strain (Qiu et al., 2013). The first target fragment, a 3.3 kb cassette containing *gpdA*-p50^Δ17-25^ was assembled via fusion PCR. This process began by amplifying two constituent fragments—N-2.4 kb and C-0.9 kb—from JZ1168 (gpdA-p50-GFP) genomic DNA using primers P50NdNC (5’-GACGGTGGAGGCTTCGTCTGTAGCGAGATCCTGGGCGAGT-3’) /P50NN (5’-AGGGAGGTTTGAACCATGG-3’) and P50NdCN (5’- ACAGACGAAGCCTCCACCG-3’) /P50CC (5’- AAGTGCTTCAGCGTCTGCTG -3’). The complete 3.3 kb fragment was then synthesized using the internal primers p50NN1(5’-TCGGAGATGGTTCGATCCTG -3’) and p50CC1(5’-TTCCTGACGCGGGTAGAAAG -3’). Concurrently, a second, 4.0 kb S-p50-ApyrG fusion fragment was generated by PCR amplification from the SX1 strain, utilizing the P50GNN (5’-AAGATGAGATGGCGGCGTC -3’) and P50GCC (5’- TCCAACCACACAAGGAATG -3’) primer set. Following co-transformation of these two fragments, successful transformants were screened for high p50^Δ17-25^-S expression via western blots and verified through PCR and sequencing using the gpdAC5 (5’-CAGTTCGAGCTTTCCCACTT-3’) and AfpyrGN3 (5’-CGAAGAGGGTGAAGAGCATT-3’) primers.

### Construction of the *gpdA*-p50-GFP strain

The gpdA promoter fragment (1.23kb) was amplified via PCR using genomic DNA from strain RQ247 as a template and the primer pair GPDF (5’-GACTCGAGTACCATTTAATTCTATTTG-3’)/GPDR (5’- TGTGATGTCTGCTCAAGCG -3’). Simultaneously, the N-terminal and C-terminal fragments of p50 (each 1kb) were amplified from GR5 genomic DNA using primer sets P50NN (5’-AGGGAGGTTTGAACCATGG-3’) /gpdFp50NC (5’-CAAATAGAATTAAATGGTACTCGAGTCAGCTAGAATATTGAAGGATCTTAGTTGTC-3’), and p50CC (5’- AAGTGCTTCAGCGTCTGCTG -3’) /gpdRp50ATG (5’-CCCGCTTGAGCAGACATCACAATGGCTTTCAACAAAAAATATGCTGGTC-3’), respectively. These three fragments were subsequently fused by overlap extension PCR using primers p50NN1 and p50CC1 (5’- TTCCTGACGCGGGTAGAAAG -3’) to generate a 3.3 kb gpdA-p50 cassette. A 4.7 kb p50-GFP-AfpyrG fusion fragment was then amplified from the genomic DNA of strain SX2 (Zhang et al. 2024) using primers p50GNN1 (5’- CGAGGCGAAGGACACATCA - 3’) and p50GCC1 (5’- CATCTTTAACAGCTGCCGCC -3’). Finally, the 3.3 kb gpdA-p50 fragment and the 4.7 kb p50-GFP-AfpyrG fragment were co-transformed into the XY42 strain. Transformants exhibiting GFP signals under fluorescence microscopy were isolated and verified by PCR and sequencing using primers: gpdAC5 (5’-CAGTTCGAGCTTTCCCACTT-3’) and GFP5R (5’-CCAGTGAAAAGTTCTTCTCCTTTAC-3’).

### Construction of a *gpdA*-p50^Δ17-25^-GFP strain

The construction of the *gpdA*-p50^Δ17-25^-GFP strain involved the co-transformation of two distinct PCR products into the XY42 host strain (Zhang, et al. 2024). The first target fragment, a 3.3 kb cassette containing the gpdA-p50 N terminus 17-25 amino acids deletion, was assembled via fusion PCR. This process began by amplifying two constituent fragments—N-2.4 kb and C-0.9 kb—from JZ1168 (gpdA-p50-GFP) genomic DNA using primers P50NdNC (5’-GACGGTGGAGGCTTCGTCTGTAGCGAGATCCTGGGCGAGT-3’) /P50NN (5’-AGGGAGGTTTGAACCATGG-3’) and P50NdCN (5’- ACAGACGAAGCCTCCACCG-3’) /P50CC (5’- AAGTGCTTCAGCGTCTGCTG -3’). The complete 3.3 kb fragment was then synthesized using the internal primers p50NN1(5’-TCGGAGATGGTTCGATCCTG -3’) and p50CC1(5’-TTCCTGACGCGGGTAGAAAG -3’). Concurrently, a second, 4.7 kb p50-GFP-ApyrG fusion fragment was generated by PCR amplification from the SX2 strain, utilizing the P50GNN (5’-AAGATGAGATGGCGGCGTC -3’) and P50GCC (5’- TCCAACCACACAAGGAATG -3’) primer set. Following co-transformation of these two fragments, successful transformants were screened for high p50-GFP expression via western blots and conclusively verified through PCR and sequencing using the gpdAC5 (5’-CAGTTCGAGCTTTCCCACTT-3’) and GFP5R (5’-CCAGTGAAAAGTTCTTCTCCTTTAC -3’) primers.

### Construction of the p50-mCherry strain

To construct the p50-mCherry fusion at the endogenous *p50* locus, a 4.7 kb *p50-*mCherry*-AfpyrG* cassette was generated using overlap extension PCR. First, three distinct fragments were amplified: a 2.7 kb mCherry*-AfpyrG* fragment was amplified from the genomic DNA of strain RQ413 using primers GAGAF (5’- GGAGCTGGTGCAGGCGCTG-3’) and pyrG3 (5’-CTGTCTGAGAGGAGGCACTGAT-3’); a 1 kb N-terminal flanking fragment and a 1 kb C-terminal flanking fragment were amplified from wild-type genomic DNA using primer pairs p50GNN (5’- AAGATGAGATGGCGGCGTC-3’), /p50GNC (5’-GCTCCAGCGCCTGCACCAGCTCCCTTCCCACTCTCCAACTTCTCC-3’), and p50GCN (5’-CATCAGTGCCTCCTCTCAGACAGAGTACTTAATATAGTGTAAGGTGAGATG-3’), /p50GCC (5’-TCCAACCACACAAGGAATG-3’), respectively.

These three fragments were subsequently fused by PCR using primers p50GNN1 and p50GCC1 to produce the final 4.7 kb transformation cassette. The resulting fragment was transformed into the TNO2A3 (Δku70) strain (Nayak et al., 2006). Transformants were screened for mCherry expression via fluorescence microscopy. Homologous integration at the p50 locus was confirmed by PCR amplification using primers AfpyrG5 (5’-AGCAAAGTGGACTGATAGC-3’) and p50GCC2 (5’- GTCCAGATATATGACGTACCTAGCG -3’).

### Construction of the Arp11-mCherry strain

To construct the Arp11-mCherry fusion at the Arp11 locus, three initial fragments were amplified. A 2.7 kb *mCherry-AfpyrG* fragment was amplified from the genomic DNA of strain RQ413 using primers GAGAF (5’- GGAGCTGGTGCAGGCGCTG -3’) and pyrG3 (5’- CTGTCTGAGAGGAGGCACTGAT-3’). Two additional 1 kb fragments, representing the N-terminal and C-terminal flanking regions of the integration site, were amplified from wild-type GR5 genomic DNA using primer pairs A11GNN (5’- TGTTTACGCGGCTTTTGGGC -3’), /A11GNC (5’- GGCTCCAGCGCCTGCACCAGCTCCCCCCCACCCTGCCAACGTCC -3’), and A11GCN (5’- ATCAGTGCCTCCTCTCAGACAGTTAGCCGGGAGGCTTTTCTC -3’) /A11GCC (5’-TTCCCTGGCGACGTCCTCAC-3’), respectively. These three fragments were subsequently fused via overlap extension PCR using primers A11GNN and A11GCC to generate a final 4.7 kb Arp11-*mCherry-AfpyrG* cassette. This cassette was transformed into the TNO2A3 (τι*ku*70) strain. Transformants were screened for mCherry expression using fluorescence microscopy, and successful homologous integration was confirmed by PCR using primers AfpyrG5 (5’-AGCAAAGTGGACTGATAGC-3’) and A11GCC2 (5’- CATGCGCTACAGGGCTTCC-3’).

### Biochemical pull-down assays, western analysis and antibodies

To harvest cells, overnight grown fungal mycelia (or hyphal mass) were filtered through miracloth (EMD Millipore Corp.) and washed with ∼200 ml distilled water. Excess liquid was removed by pressing the folded miracloth (containing the hyphal mass inside) between paper towels. After the liquid has been removed, the weight of the hyphal mass (without the miracloth) was measured on a balance, and about 0.6 g hyphal mass was collected and wrapped with a foil paper, labeled and stored in a -80°C freezer. To break hyphal cells, a mortar and pestle were used to grind the frozen 0.6 g hyphal mass to a fine powder in liquid nitrogen (Basically, liquid nitrogen was added into the mortar a little bit at a time during grinding just to keep the sample frozen). Cell extracts were prepared adding 1.5 ml of lysis buffer containing 50 mM Tris-HCl (pH 8.0), 0.01% Triton X100 and 10 μg/mL of a protease inhibitor cocktail (Sigma-Aldrich). Cell extracts were centrifuged at 20,000 *g* for 60 minutes at 4°C, and supernatant was used for the pull-down experiment.

The μMACS GFP-tagged protein isolation kit (Miltenyi Biotec) was used to pull down proteins associated with the GFP-tagged protein, which was done as described previously (Zhang et al., 2014). To pull down GFP-tagged proteins, 35 μL anti-GFP MicroBeads were added into the cell extracts for each sample and incubated at 4°C for 60 minutes. The MicroBeads/cell extracts mixture was then applied to the μColumn followed by gentle wash with 200 μL of the same lysis buffer used above for protein extraction. After washing three times, 80 μL of pre-heated (95°C) 2 x SDS-PAGE sample buffer was used to elute the proteins. The eluted proteins were applied to premade 4–15% Criterion™ TGX Stain-Free™ Protein Gels (Bio-Rad), and after finishing running, proteins in the gels were transferred overnight at 4°C to nitrocellulose membrane (0.45 μm) (Bio-Rad). Western blot analyses were performed using the alkaline phosphatase system and blots were developed using the AP color development reagents from Bio-Rad. Quantitation of the protein band intensity was done using the Image Studio Lite software (version 5.2). Specifically, an area containing the whole band was selected as a region of interest, and the intensity sum within the region of interest was measured. Then, the region of interest box was dragged to the equivalent region of the negative control lane or a blank region without any band on the same blot to take the background value, which was then subtracted from the intensity sum. The rabbit polyclonal antibody against GFP was from Takara Bio Inc (Catalog number: 632592). The rabbit monoclonal antibody against the S-tag was from Cell Signaling Technology (Catalog number: 12774S). Polyclonal antibody against Arp1 was generated using the service of Pacific Immunology (www.pacificimmunology.com). An immunograde peptide of Cys-SADEWHEDPEIIHRKFA from the C-terminus of Arp1 was synthesized and conjugated to the KLH carrier protein, and this conjugated form was used as an antigen for rabbit injection. The antibody was affinity-purified using the same peptide. Similarly, polyclonal antibody against p150 was generated using the service of Pacific Immunology (www.pacificimmunology.com), and an immunograde peptide of Cys-PDRKANAVQPVEPAIEPTF from the C-terminus of p150 was used. Polyclonal antibody against Arp11 was generated using the service of Boster Antibody and Elisa experts (www.bosterbio.com). A recombinant protein of 217 amino acids (RSALVVDIGWAETVVSGIYEYREVTTKRSTRAMRSLIQETGRMFTRLLGGDSQPDTISVEFEFCEEVVSRFAWCQPSRSGYYKETAENSLADILDKTISIPSPSNPGSSDIELPFSKLEELVEKVLLAQGMADSDLDDQEKPISLLVYNTLLSLPPDVRGICMSRIVFVGGGANIAGIRSRILDEVAHLIELYGWSPVRGRLIEQQIQKLQSLKLSQ) was used for rabbit injection and purification of serum.

### Statistical analysis

All statistical analyses were done using GraphPad Prism 10 for macOS (version 10.6.1, 2025). The D’Agostino & Pearson normality test was performed on all data sets except for data sets with small n (n=3). For western blot data, data distribution was assumed to be normal but this was not formally tested because the n number is 3 for these data sets. For these data, Welch’s *t*-test (unpaired, two-tailed) was used for analyzing any two data sets and Ordinary one-way ANOVA with Dunnett’s multiple comparison test was used for analyzing any three or four data sets. For all other data sets, the D’Agostino & Pearson normality test was performed. The data sets that did not pass the D’Agostino & Pearson normality test (alpha=0.05) were analyzed using the Mann-Whitney test (unpaired, two tailed), a non-parametric test without assuming Gaussian distribution. The data set that passed the D’Agostino & Pearson normality test were analyzed using the student *t*-test (unpaired, two tailed).

### AlphaFold3-based structural prediction

Protein three-dimensional structures were predicted using AlphaFold 3 (accessed via the AlphaFold Server, https://alphafoldserver.com) in template-free mode. Predicted models were subsequently inspected and analyzed using UCSF ChimeraX.

## Acknowledgements

We thank Andrew Carter and Erika Holzbaur for helpful discussions, and Andrew Carter for sharing structural predictions that stimulated experiments and for helpful comments on the manuscript. This work was funded by the National Institutes of Health R35GM140792 (to X.X.) and a Department of War #OSD(HA).2022ICD.WBH-1 Joint DOTML-PF Warfighter Brain Health grant to USUHS Department of Biochemistry and Molecular Biology. Disclaimer: The opinions and assertions expressed herein are those of the author(s) and do not necessarily reflect the official policy or position of the Uniformed Services University or the Department of War.

---

**Table S1.**
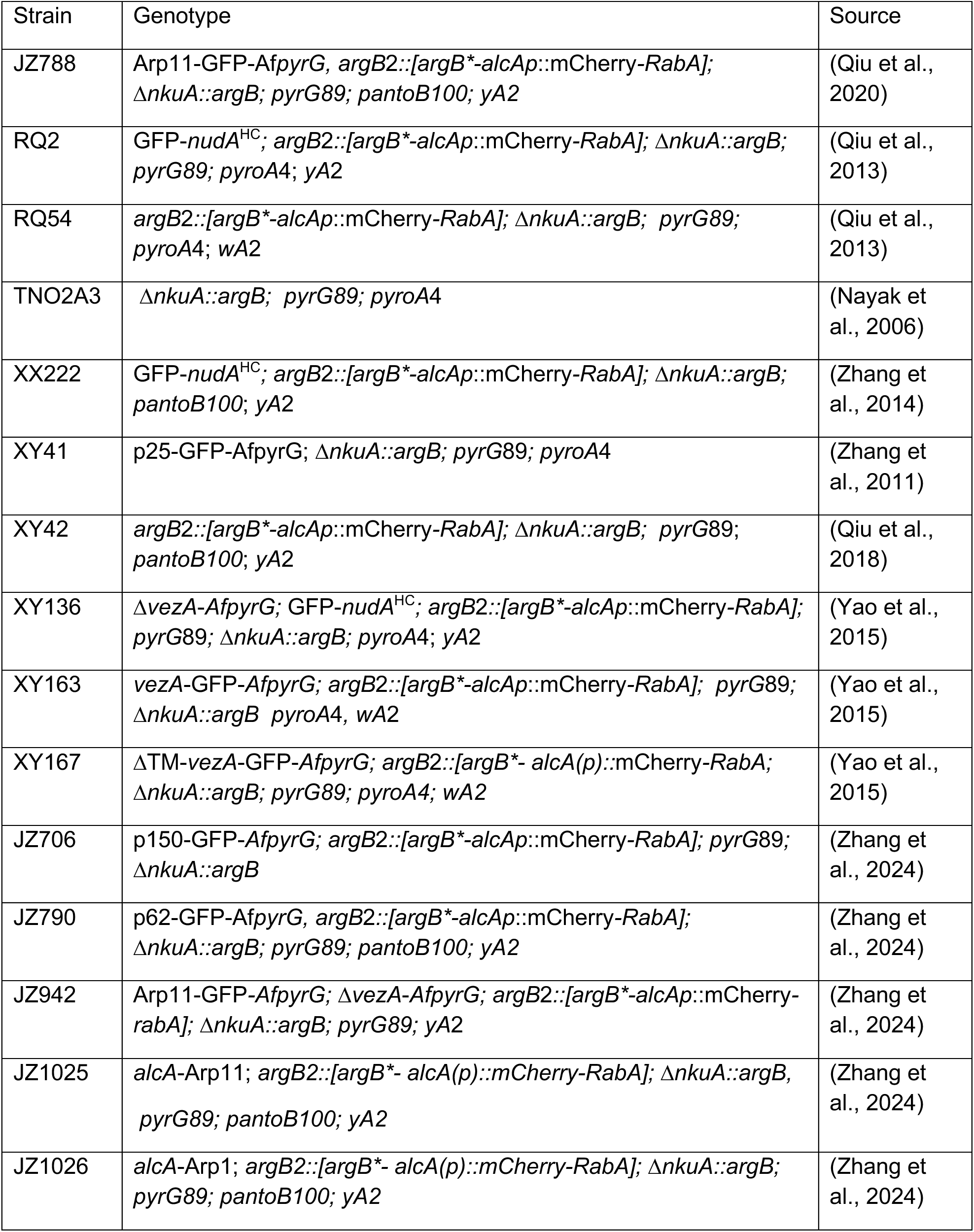
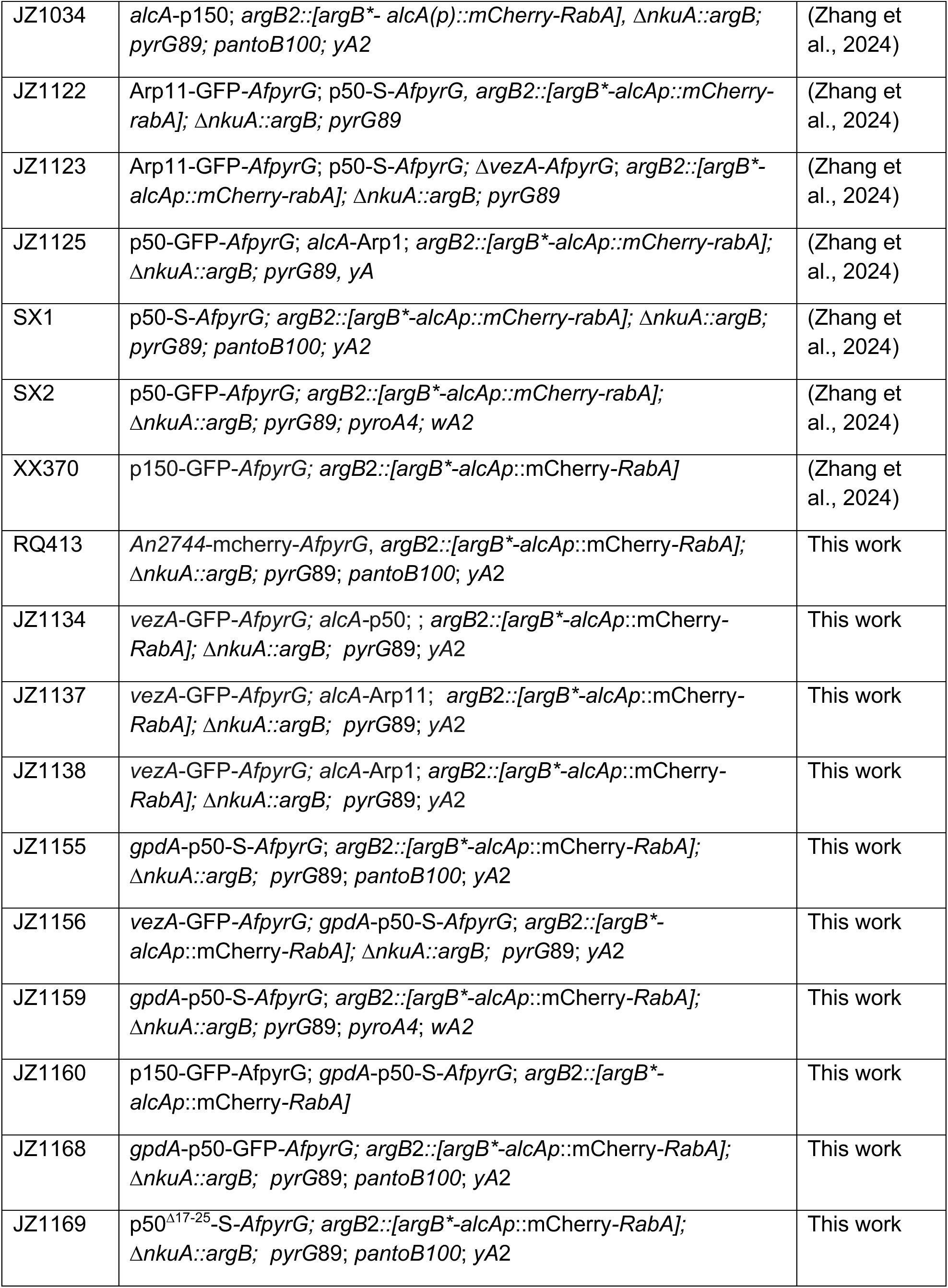
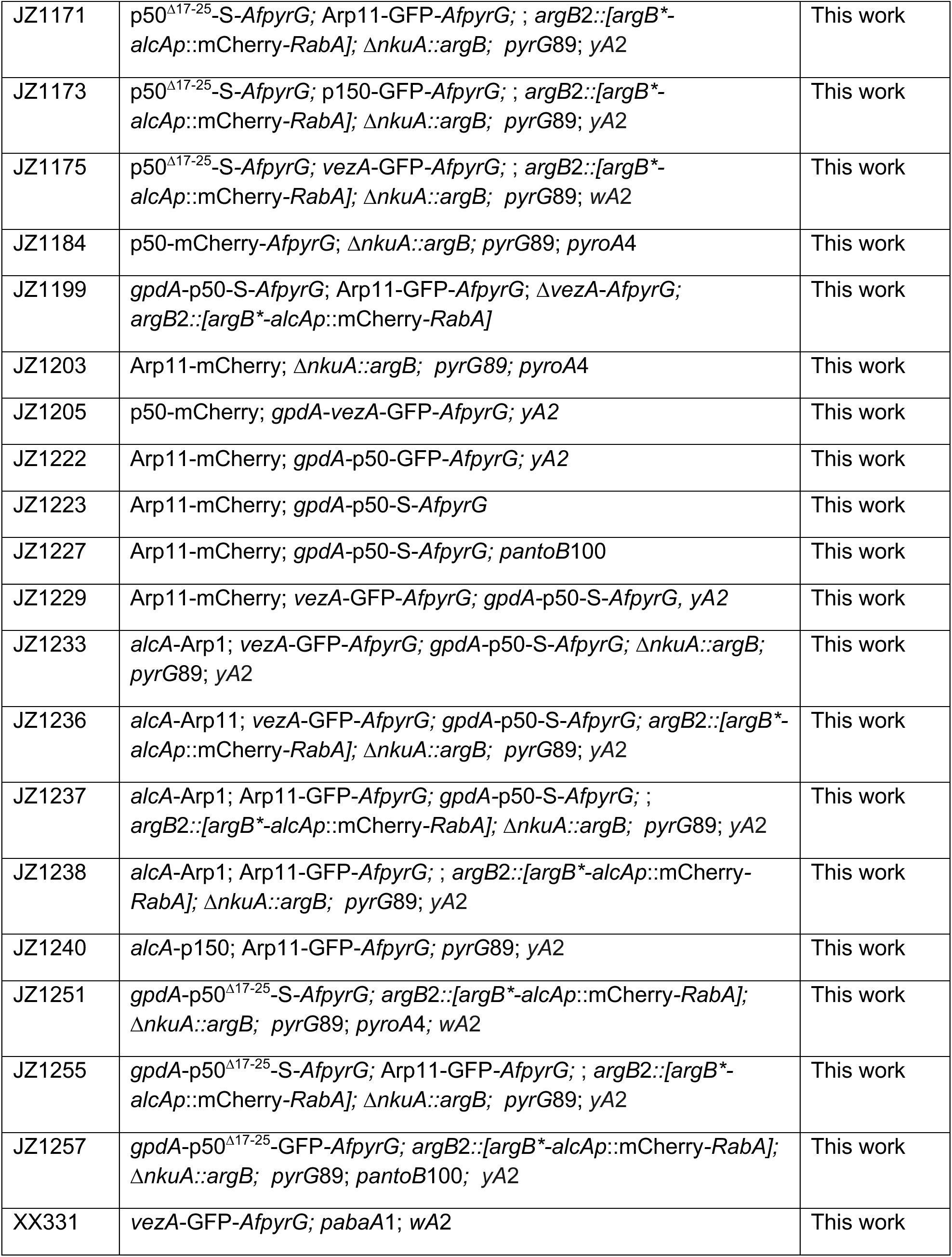
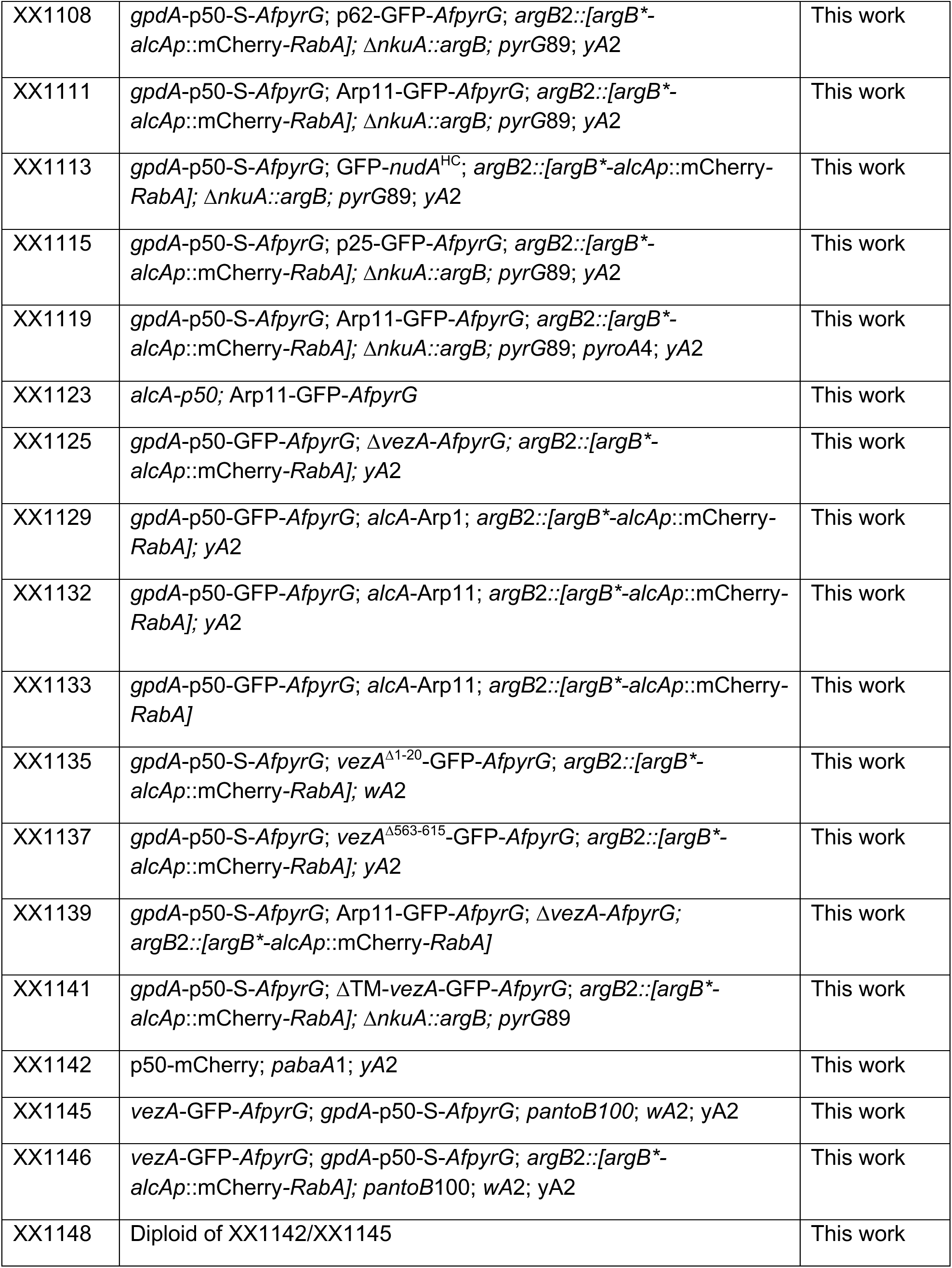
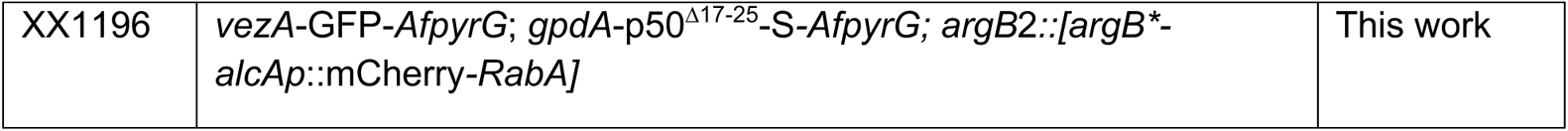
***Aspergillus nidulans* strains used in this study**

**Figure S1.**
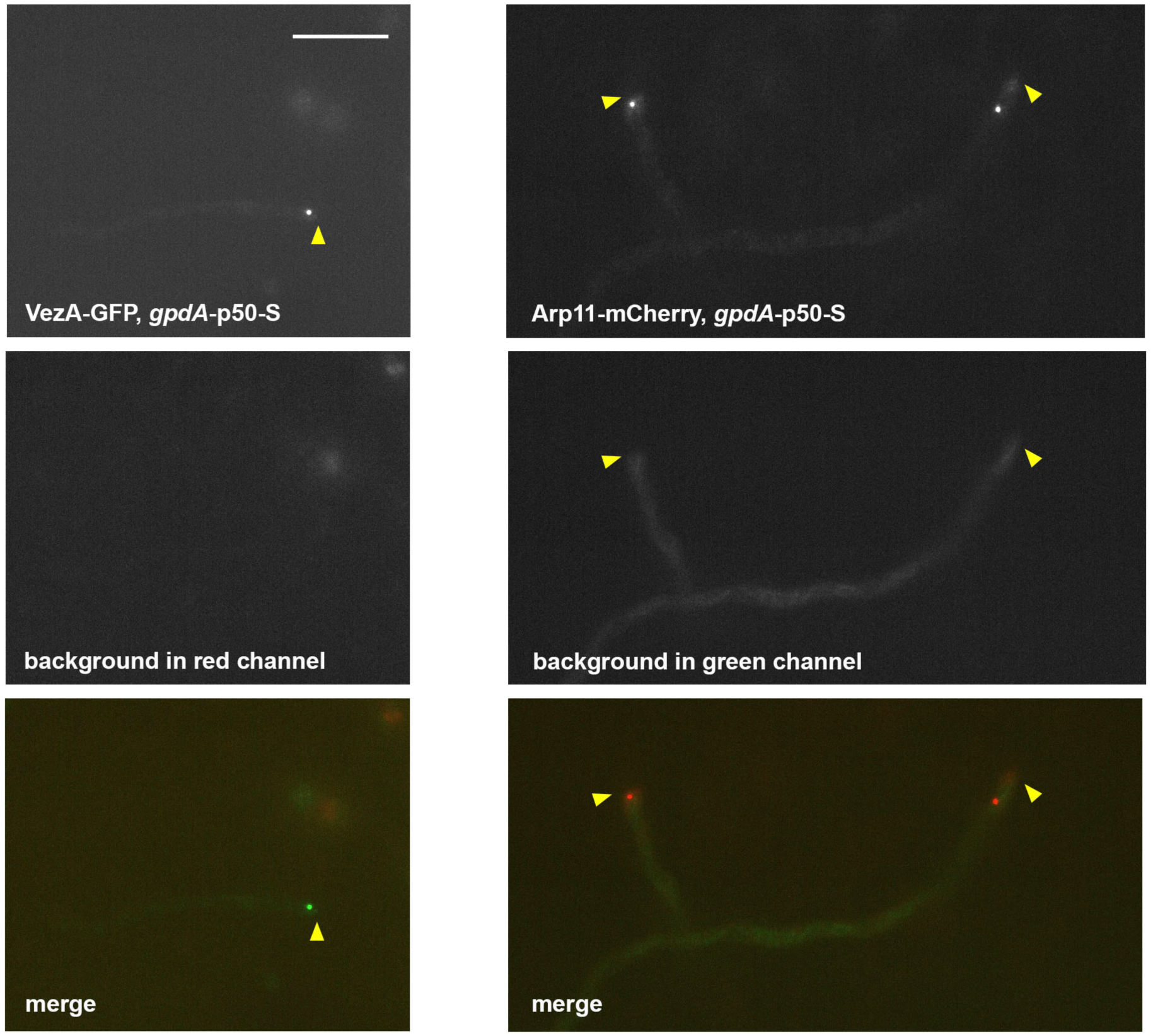
Controls for the images from the strain containing Arp11-mCherry, VezA-GFP and *gpdA*-p50-S (see Figure 4E middle panels). The purpose of these controls is to show that the clustered VezA-GFP signals can be seen only in the green channel and the clustered Arp11-mCherry signals can be seen only in the red channel under our imaging conditions. Images of a strain containing VezA-GFP and *gpdA*-p50-S are shown in the left panels, and those of a strain containing Arp11-mCherry and *gpdA*-p50-S are shown in the right panels. Hyphal tip is indicated by a yellow arrowhead. Bar, 10 μm.

**Figure S2.**
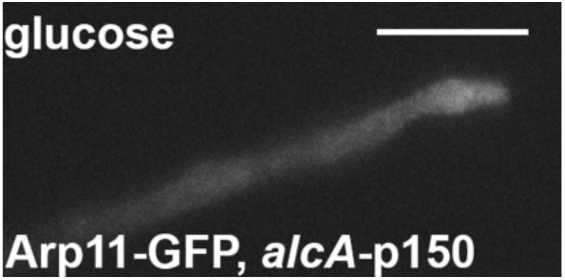
Arp11-GFP does not form clusters upon loss of p150. In the *alcA*-p150 background where the expression of p150 is repressed on glucose medium, Arp11-GFP signals are diffuse and slightly enriched at the hyphal tip but does not form clusters, which is in contrast to the cluster formation of Arp11-GFP in the *gpdA*-p50-S background (see Figure 2A and Figure 6A). Bar, 10 μm.

**Figure S3.**
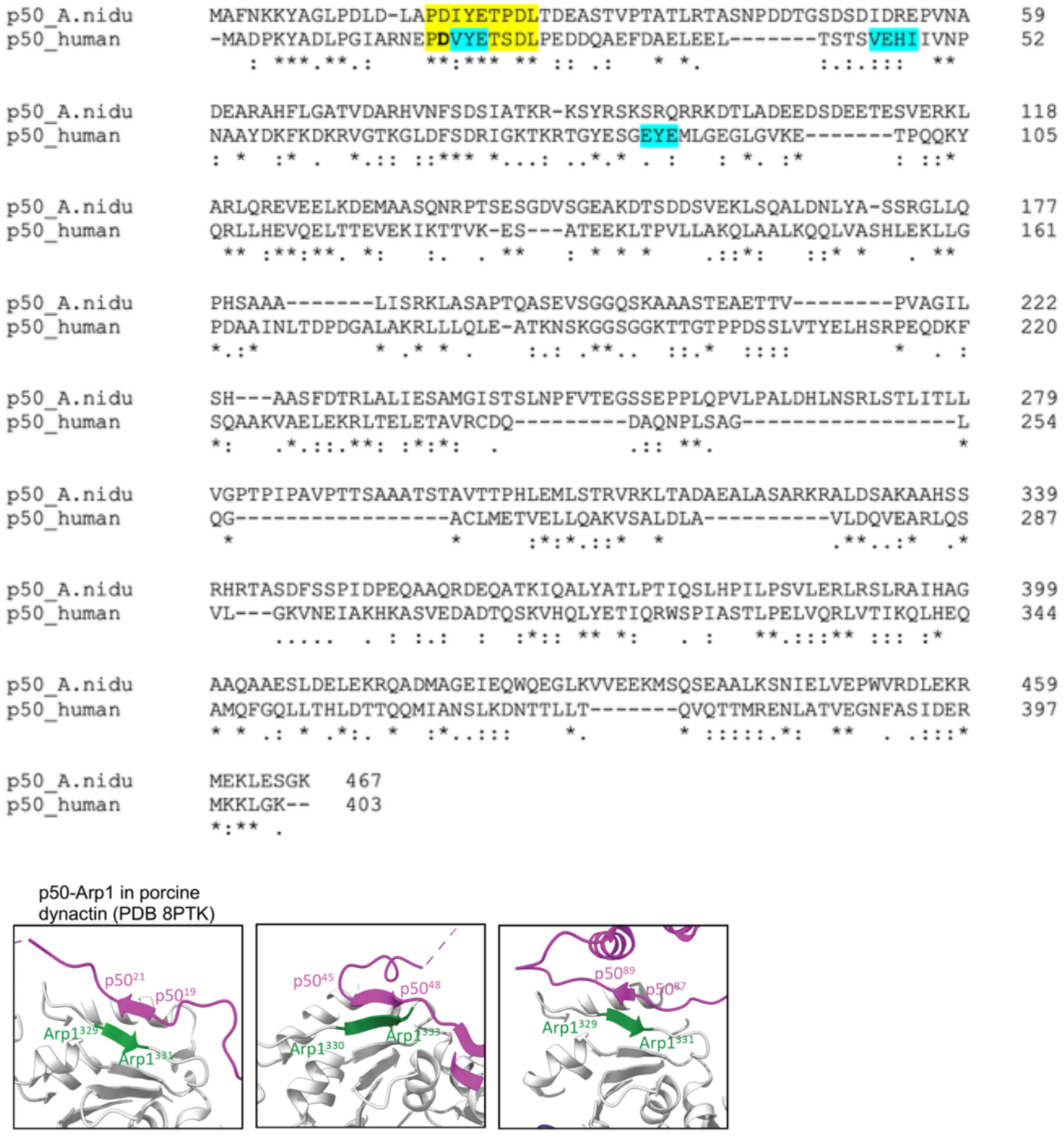
Protein sequence alignment of p50 proteins and beta-sheet structures of p50-Arp1 interaction. A sequence alignment of *A. nidulans* p50 and human p50. The amino acids involved in beta-sheet structures of p50-Arp1 interaction are highlighted by the cyan background. The p50 amino acids 17-25 (in both *A. nidulans* and human) around the first beta-sheet structure are highly conserved (yellow background). The alignment was done using the CLUSTAL O(1.2.4) multiple sequence alignment program. Residues that are identical (*), strongly similar (:) or weakly similar (.) are indicated. The structures of the porcine p50-Arp1 interactions shown below the alignment are based on published structure data from PBD 8PTK (Singh et al., 2024).

